# OpenPathSampling: A Python framework for path sampling simulations. I. Basics

**DOI:** 10.1101/351494

**Authors:** David W.H. Swenson, Jan-Hendrik Prinz, Frank Noe, John D. Chodera, Peter G. Bolhuis

**Affiliations:** van’t Hoff Institute for Molecular Sciences, University of Amsterdam, PO Box 94157, 1090 GD Amsterdam, The Netherlands; Computational and Systems Biology Program, Sloan Kettering Institute, Memorial Sloan Kettering Cancer Center, New York, NY 10065, USA; Department of Mathematics and Computer Science, Arnimallee 6, Freie Universität Berlin, 14195 Berlin, Germany

**Keywords:** transition path sampling (TPS), transition interface sampling (TIS), molecular dynamics simulation (MD), rare events

## Abstract

Transition path sampling techniques allow molecular dynamics simulations of complex systems to focus on rare *dynamical events*, providing insight into mechanisms and the ability to calculate rates inaccessible by ordinary dynamics simulations. While path sampling algorithms are conceptually as simple as importancesampling Monte Carlo, the technical complexity of their implementation has kept these techniques out of reach of the broad community. Here, we introduce an easy-to-use Python framework called OpenPathSampling (OPS) that facilitates path sampling for (bio)molecular systems with minimal effort and yetis still extensible. Interfaces to OpenMM and an internal dynamics engine for simple models are provided in the initial release, but new molecular simulation packages can easily be added. Multiple ready-to-use transition path sampling methodologies are implemented, including standard transition path sampling (TPS)between reactant and product states, transition interface sampling (TIS) and its replica exchange variant(RETIS), as well as recent multistate and multiset extensions of transition interface sampling (MSTIS, MISTIS).In addition, tools are provided to facilitate the implementation of new path sampling schemes built on basic path sampling components. In this paper, we give an overview of the design of this framework and illustrate the simplicity of applying the available path sampling algorithms to a variety of benchmark problems.

## I. INTRODUCTION

Biomolecular systems, such as proteins and nucleic acids, can undergo complex conformational changes on long timescales that are challenging for atomistic molecular simulations to reach. Forexample, atomistic moleculardynamics (MD) must employ timesteps on the scale of femtoseconds to faithfully reproducethefastestvibrational modesto maintain simulation stability and fidelity, while the kinetic timescales (e.g. of protein folding or binding) can often range from microseconds to seconds or more. In protein-ligand binding, mean residence times for bound druglike molecules are often several hours, presenting an enormous challenge to studying dissociation mechanisms or predicting unbinding rates by straightforward MD [1–3]. In these and othersituations, simulating a sufficient number of these rare events (folding/unfolding or binding/unbinding) to produce a statistically meaningful description of the dominant mechanism or estimate of rate constants is often so challenging as to be untenable by straightforward means. Slow kinetic time scales primarily arisefrom largekineticbarriers between metastablestates[4–7]. The observed dynamics is dominated by long waiting times within metastable basins, punctuated by rare events of interest occurring over a short time [8]. Straightforward molecular simulation is highly inefficient as most effort will be wasted simulating uninteresting dynamics as the system remains trapped within metastable states [9].

One approach to overcoming the rare event problem is to bias the potential energy surface or alterthe probability density of sampled conformations to enhance the occurrence of the rare event. *A priori* knowledge of a suitable reaction coordinate allows the use of biasing potentials or higher effective temperatures, reducing effective free energy barriers. Many such enhanced sampling methods have been developed (e.g. see Refs. [10–20]). Useful bias potentials capable of enhancing the frequency of rare events require (a set of) collective variables that approximate the reaction coordinate; poor choices will lead to poor sampling of the reactive pathways, and hence poor estimates of the dynamical bottlenecks and the related barrier heights and rates. Even worse, some methods are sensitive to the omission of slow degrees of freedom, and may lead to incorrect models of the reactive pathways. In general, removing the effect of the bias potential to yield correct dynamics is difficult.

*Path sampling techniques*, in paticular *transition path sampling* [9, 21–23], provide a solution to the rare event problem without requiring the same degree of knowledge of reactive pathways. Instead of biasing the potential-which leads to heavily perturbed dynamics-these techniques bias the probability with which a given transition path is sampled, without perturbing these paths themselves. This property allows the unbiased equilibrium dynamics to be recovered. For the simple case of a two-state system separated by a single barrier, the straightforward MD simulation time to observe a number of transitions scales exponentially in the barrier height. In contrast, transition path sampling only focuses on short parts of the MD trajectory that traverse the barrier, providing exponential acceleration in the sampling of rare events [9,22]. Other methods based on trajectory sampling include forward flux sampling (FFS) [24], adaptive multilevel splitting [25], milestoning [26], the RESTART methodology [27],SPRESS [28], NEUS [29], Weighted Ensemble [30,31] and many others.

In addition to studying rare events directly, path sampling methods can be combined with other approaches for describing statistical conformational dynamics. For example, Markov state models (MSMs) have emerged as a popular way to represent the longtime statistical dynamics of complex processes involving many distinct metastable conformational states [32]. By discretizing conformation space and describingstochastictransitions between regions with a transition or rate matrix, MSMs can describe the long-time statistical dynamics of complex systems with bounded approximation error [32]. While standard MSM construction approaches utilize large quantities of unbiased simulation data, path sampling techniques can be utilized to rapidly construct or improve MSM transition matrices by focusing on harvesting trajectories for poorly sampled transitions [2,33–37]. More recently, techniques have emerged for combining both biased and unbiased dynamics to construct *multi-ensemble Markov Models (MEMMs)* [38–41], enabling even richer combinations of multiple efficient sampling techniques for rapid construction of statistical models of dynamics.

While transition path sampling techniques are very flexible, the complexity of their implementation and lack of a standard tool for applying them has slowed their adoption. In particular, many path sampling techniques require monitoring of dynamics to detect when stopping conditions are reached, and the control of and integration with standard simulation packages has been a practical obstacle for widespread use. As a solution to this, we have developed a new framework called **OpenPathSampling (OPS)** that enables path sampling techniques to be employed in a flexible, general manner. This framework is “batteries included”, with a number of different path sampling algorithms and worked examples available that can help users to apply path sampling techniques on their own system. Both low-dimensional toy model systems and complex molecularsystems are supported, with complex systems supported using interfaces to external simulation codes. Currently, OPS supports the GPU-accelerated molecularsimulation code OpenMM [42,43], although support for other codes can be added. The framework is flexible and extensible, allowing users to easily explore implementation of new path sampling algorithms in addition to applying or extending existing algorithms or connecting new simulation codes. Many other methods, such as FFS or mile-stoning, could also be implemented within the framework of OPS. For the sake of clarity, however, we will limit ourselves here to the transition path sampling based methods [^1^ Note that in this work we often use ‘transition path sampling’ and ‘path sampling’ interchangeably. The reason is that the concept of path sampling is more inclusive, and also covers algorithms that do not immediately aim to cross (single) barriers. However, it is understood that all path samplingmethods in this work fall into the larger ‘transition path sampling’ family of algorithms.]. OPSdif-fers in scope and versatility from the PyRETIS package[44], a recently developed package to conduct advanced transition path sampling simulations.

In this paper, we first give a brief overview of a variety of path sampling techniques that are implemented in the OPS framework (Section II); explain how the basic path sampling concepts relate to OPS object classes (Section IV); review the general workflow associated with setting up, running, and analyzing a path sampling calculation (Section V); and then provide a number of detailed examples that illustrate the flexibility and simplicity of applying various path sampling techniques using this framework (Section VI). In the process of developing a framework capable of easily implementing a multitude of path sampling techniques, we have significantly generalized the manner in which path ensembles can be constructed and used within the path sampling mathematical framework. While this expressive path ensemble specification language is briefly introduced (Section III) and utilized in the examples described here, this approach is described in detail in a companion paper in this issue [45].

## II. BACKGROUND

### A. The concept of path ensembles

Here, we presume the reader is somewhat familiar with the transition path sampling literature [9, 21–23, 46]. While we give a brief overview of the main concepts in this section, readers not familiar with this topic are encouraged to start with a basic review such as Ref. [46].

The types of path sampling considered in this paper—and implemented and supported by OpenPathSampling—deal with equilibrium dynamics, obeying microscopic reversibility, so that the stationary distribution is the Boltzmann distribution. Moreover, *ergodicity* is assumed; that is, an infinitely-long trajectory has a nonzero probability to visit every point in phase space. This guarantees that (dynamical) averages computed in the path ensemble, such as rate constants, are identical to those of an infinitely long trajectory.

A *path* or *trajectory* consists of a sequence of *L* + 1 points in configuration or phase space x ≡ {*x*_0_, *x*_1_,…,*x*_*L*_} generated by some dynamical model (such as Hamiltonian, Langevin, Brownian, or even Monte Carlo dynamics), with the initial configuration *x*_0_ drawn from an initial (equilibrium) distribution ρ(*x*_0_). The *path ensemble* is defined by the probability distribution 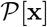 of such paths (with the length *L* either fixed or varying), and can be sampled using a Markov Chain Monte Carlo (MCMC) algorithm. Path sampling algorithms consist of a few main ingredients: (1) a scheme for initializing the sampler with an initial path; (2) one or more schemes for proposing new trial paths from the current path; (3) an acceptance criteria (e.g., based on Metropolis-Hastings) used to accept or reject the proposed trial path to generate a new sample from the path probability density (ensemble) of interest.

The idea of path sampling is to enhance the probability sampling of certain paths, either by biasing the path probability or by constraining the path ensemble. Analogous to how standard Monte Carlo importance sampling techniques can enhance sampling of rare configurations by multiplying the probability density by a biasing factor *w*_*bias*_(*x*) based on the instantaneous conformtion *x*,

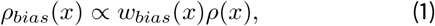

and subsequently using trns Dias to unbias me sampled ensemble and recover equilibrium expectations, path sampling techniques can enhance the sampling of rare trajectories by multiplying by a biasing weight *w*_*bias*_ [x], based on the trajectory x,

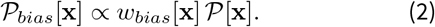

Many types of path sampling, notably standard *transition path sampling* (TPS) [9,21], define constrained path ensembles which select trajectories that begin in one region of configuration space A and end in another region B. OPS supports a simple but powerful way of defining path ensembles, described briefly in Section V B and expanded upon in detail in a companion paper [45]. Below, we give a brief overview of common kinds of transition path sampling simulations supported by OPS.

### B. Transition path sampling

The *transition path sampling* (TPS) [9,21] method attempts to harvest trajectories connecting two specific regions of configuration space, such as a reactant and product separated by a single free energy barrier. The constrained path ensemble for a fixed length *L* is thus

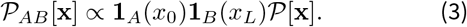

Here, x ≡ {*x*_0_, *x*_1_,…, *x*_*L*_} is a discrete-time trajectory of snapshots, 1_A_(*x*_0_) and 1_*B*_ (*x*_*L*_) are indicator function that are unity if the trajectory starts with *x*_0_ ∈ *A* and ends with *x*_*L*_ ∈ *B* and zero otherwise, and 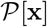 is the equilibrium path probability density. In a TPS simulation, new trial trajectories are proposed from the current sampled trajectory by selecting a phase space point along the trajectory, applying a perturbation (usually of the momenta), and “shooting” forward and backward by integrating the equations of motion until a trajectory of the original length is generated. The trial trajectory is then accepted or rejected with a Metropolis-Hastings criterion. For the simplest case of drawing the *shooting point* uniformly from the current trajectory, assigning a new velocity from the Maxwell-Boltzmann distribution, and imposing the trajectory of fixed length to begin in state *A* and end in *B*, this acceptance criteria amounts to accepting the new trajectory when it satisfies the defined ensemble of interest by terminating in regions *A* and *B*; the old path is otherwise retained if the proposed trajectory is rejected. Depending on the details of the shooting move, the exact acceptance criteria will take on different forms [9,21–23,46].

Transition path sampling is immensely powerful, as the difficult problem of describing reaction mechanisms is reduced to the much easier problem of defining stable states *A* and *B*. Reactive trajectories are efficiently harvested because the trial trajectory quickly decorrelates from the original trajectory, yet is still likely to meet the same path ensemble constraints, such as connecting the reactant and product regions of configuration space *A* and *B*.

In order for the reactive trajectories connecting metastable sets *A* and *B* to be useful for computing transition rates and physical interpretation of mechanisms, the system must commit to and remain in the metastable states fora longtime after encountering them, i.e., transitions between *A* and *B* are rare events on the molecular time scale. The states *A* and *B* are generally defined as configurational space regions within the basin of attraction of the distinct metastable states. Trajectories initiated from configurations in these regions, called *core sets*, should have a high probability (close to unity) to remain in or quickly return to the coreset rather than escape to other states, even at the boundary of these sets [32,47].

TPS can also be used with flexible-length trajectories that are constrained to terminate when they encounterthe boundary of core sets A and B. This can be encoded in the path ensemble definition by demanding that frames 1 to *L*-1 are neither in A nor in B. This approach is more efficient at sampling reactive trajectories by avoiding sampling long dwell times in each state at either end of the trajectory [48]. To maintain detailed balance, the acceptance criterion then contains the ratio of the previous and trial path length, i.e., the number of frames from which the shooting point is randomly chosen. TPS can also easily be extended to multiple states by allowing more states in the path ensemble definition [34]. A variety of other path proposal moves have been described to attempt to increase acceptance probabilities in certain regimes, including shifting moves [21], small velocity perturbations [21], precision shooting [49], permutation shooting [50], aimless shooting [50], and spring shooting [51].

### C. Transition interface sampling (TIS)

While TPS yields information about the mechanism of the rare events, important quantities such as the kinetic rate constant requires an additional scaling factor that quantifies how frequent transition paths are relative to non-transition paths. Therefore, one has to relate the constrained TPS ensemble with the unconstrained path ensemble, as given by an infinitely long ergodic unbiased MD trajectory [9]. This unconstrained *total* (or complete) path ensemble comprises the set of path ensembles starting from each stable state, consisting of all (properly weighted) paths that leave that state and either return to it, or go on to any other stable state. Even when restricting the path ensemble to start in a partic-ularstate A, straightforward path sampling of an otherwise unconstrained ensemble is naturally very inefficient, as the important transitions to other states are exceedingly rare. However one can construct the total path ensemble (for each state) by a staging procedure. In such a procedure one can constrain the paths to reach further and further out of the state (while of course still starting in the stable state). This constraining can be done using the transition interface sampling method (TIS) [52], an extension of TPS that is explained below. Reweighting of the resulting paths yield then (an estimate of) the total path ensemble [53].

Transition interface sampling (TIS) [52] provides a more efficient evaluation of the rates compared to the original TPS rate constant calculation [54] by sampling each constrained interface ensemble. TIS defines a set of *N* nonintersecting hypersurfaces (the‘interfaces’) around the stable state, parametrized by a collective variable λ, and foliating, in principle, the entire configuration space (or even phase phase[52]). The rate constant from *A* to *B* is expressed as

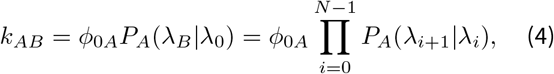

where *ϕ*_0*A*_ denotes the flux out of *A* through *λ*_0_, and *P*_*A*_(*λ*_*B*_|*λ*_0_) is the *crossing probability,* the probability that a trajectory originating from *A* reaches interface *λ*_*B*_ before returning to A, provided that the path already crosses *λ*_0_ at least once. This probability is generally low, as the transition is a rare event, but can be computed through the product of all crossing probabilitiesforthe individual interfaces, as indicated in Eq. 4, with *λ*_*N*_ ≡ *λ*_*B*_ [52]. Interfaces should be optimally placed such that each crossing probability in the product is roughly *P*_*A*_(*λ*_*i*+1_|*λ*_*i*_) ≈ 0.2 [72]. The total number of required interfacesisthusoftheorder *N* ≈ | log_5_ *P*(*λ*_*B*_|*λ*_*A*_)|. As an example, fora barrier of 30 *k*_*B*_*T*, this roughly translates into *N* = 30/ ln(5) ≈ 18 interfaces. The staging approach thus avoids the problem of the exponentially low rate in a way analogous to umbrella sampling [55].

Note that the product is not simply a product of Markovian transition probabilities, as for each interface the entire trajectory starting from *A* is taken into account. Evaluation of the crossing probabilities requires sampling the path ensemble for each interface with the constraint that the path needs to cross that interface. While trajectories could in principle be stopped when they reach the next interface, it turns out to be beneficial to continue the trajectory integration until a stable state (*A* or *B*) has been reached. This also allows the application of the so-called *reversal* move of *A*-to-*A* trajectories, where the time direction of the path is reversed, which can be done with no additional cost, but assists in decorrelating paths. The flux *ϕ*_0*A*_ can be easily obtained using straightforward MD inside state *A* [52,56].

The reverse rate can be computed by repeating the TIS simulation from state B: define a set of interfaces, sample the interface ensembles, and compute the crossing probability *P*_*B*_(*λ*_*B*_|*λ*_*A*_).

Similarto TPS, the TIS algorithm can be extended to multiple states [34]. To estimate kinetic rates between multiple states, each state *I* gets its own set of interfaces *λ*_*iI*_, and the rate constant from state *I* to state *J* is given by

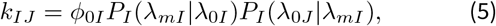

where *ϕ*_0*I*_ is again the flux from *I* through *λ*_0*I*_. The second factor is the crossing probability to an outermost interface *m*, which is typically very small and expressed as 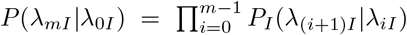. The last factor in Eq. 5 is the conditional probability that a trajectory crossing the outermost interface also reaches state *J*. The location of the outermost interfaces should be chosen such that the probability to escape from *A* is sufficiently large. Note that while interfaces belonging to state *I* constitute a foliation of non-overlapping hypersurfaces, they are completely independent from the interfaces of state *J*, and in fact are allowed to overlap [57,58].

We introduce the concept of a *transition network* [59] that, in its simplest form, represents the ensembles of paths connecting pairs of defined states. For each state in the transition network (multiple state) TIS results in a set of interface path ensembles and a straightforward MD ensemble of that stable state, which can be combined to yield the total path ensemble by reweighting. Repeating this for all states, and (again) properly reweighting [53, 60, 61], leads to an accurate description ofthe kinetic rate matrix, the free energy landscape, the mechanisms and reaction coordinates of all transitions between the metastable states. This data can be further analyzed using theory of Markovian stochastic processes, e.g., the Chapman-Kolgomorov equation [62] or transition path theory [63].

### D. Considerations in transition path sampling

The readershould beawareof a numberof challenges they may encounter in setting up transition path sampling based simulations. While an exhaustive list is beyond the scope of this paper, we list some important issues below. (See also Ref. [64]).

#### 1. Definition of the states

Transition path sampling requires knowledge ofthe stable states. Usually the stable states are easier to characterize and identify than the transition region. Analyzing straightforward MD can provide information on how to describe the states in terms of (several) collective variables. Such heuristic approaches has been used in previous applications [48, 65, 66]. In addition, tools such as clustering can be used to define the states [58]. Ideally, one would like to use automatic state recognition, and recently attempts have been made in that direction [47]. In OPS we assume that the reader has an idea about howto capture stable states by defining a range in (several) collective variables. OPS provides the user with tools to facilitate identification of these ranges, and hence definition ofthe states. The choice ofthe stable state definitions still requires careful attention, as an erroneous definition can easily lead to improper or failed path sampling. Fora detailed discussion on the stable state definitions, see Refs [22,46, 64].

#### 2. Intermediate metastable states

Even if the process of interest exhibits two-state kinetics, suggesting only two highly stable states are involved, it is possible that the presence of one or more intermediate states with lifetimes short on the overall time scale but long on the molecular timescale will cause reactive trajectories connecting the stable states to be quite long. A solution to this problem is to identify the intermediate state(s), define their coresets, and to use multistate transition interface sampling (MSTIS) [34,57]. Alternatively, one can choose to simply sample long pathways [67], which can still be quite fast given the speed of modern GPU-accelerated molecular simulation engines like OpenMM [42,43].

#### 3. Ergodicity of path space

While the TPS and TIS algorithms are “exact” in the sense that they should lead to the asymptotically unbiased estimates of path averages in the limit of infinite sampling, they suffer from the same problems that all Monte Carlo methods encounter, the problem of slowly mixing Markov chains, which in severe cases may result in broken *ergodicity* for practical computertimes. As TIS samples path space by perturbing an existing path to generate new proposals, decorrelation from the initial path to generate many effectively uncorrelated paths is essential for producing useful unbiased estimates. However, since there might be (possibly high) barriers in path space orthogonal to the interfaces between different allowed reaction channels, this is far from guaranteed. One way of solving this problem is by using replica exchange among path ensembles in transition interface sampling (RETIS) [68, 69].

### E. Replica exchange transition interface sampling (RETIS)

The RETIS algorithm simultaneously samples all TIS ensembles while allowing for swapping of paths between interface ensembles when possible [68, 69]. Atransition path that follows one particular mechanism can then slowly morph into a completely different transition path by exchanging it back and forth among all interfaces to state *B*. Including an exchange between pathways belonging to different states further enhances sampling convergence [69].

Further sampling improvement can be achieved by including van Erp’s *minus interface ensemble* [68, 69]. The minus interface move exchanges a trajectory in the first interface ensemble with a trajectory exploring the stable state (the minus interface ensemble). This serves two aims: *(1)* to decorrelate pathways in the first interface which tend to be short; and *(2)* to provide a direct estimate for the flux out ofthe stable state [68–70]. OPS includes an implementation of multiple state RETIS, which we will refer to as MSTIS.

The default MSTIS approach employs a single set of interfaces for each state, based on one order parameter. Multiple interface set TIS (MISTIS), also implemented in OPS, generalizes this approach to include multiple interface sets for states or transitions [71]. Although TIS is much less sensitive to the choice of order parameter than other enhanced sampling methods [72], in practice, the efficiency is affected by this choice. Using different order parameters to describe (sets of) interfacesfordifferenttransitionsand/orstates, with the help of replica exchange, might alleviate such efficiency problems.

A drawback ofthe (multiple state) RETIS approach is that it requires one replica to be simulated for each interface; for systems with multiple stable coresand associated interfac *𝒪*(10) interfaces. This large number of interface ensembles prevent efficient implementation ofthe method for systems more complex than toy models. A parallel implementation of all interfaces might seem a simple solution, but will be complicated by thefactthattheduration ofthe paths in the different interface ensembles varies wildly. Single replica TIS (SRTIS), based on the *method of expanded ensembles* [73], can alleviate this problem [61]. Instead of exchanging paths between interface ensembles, only one replica is sampled, and transitions between ensembles are proposed. To avoid the replica remaining close to the stable state interface ensemble, one needs a biasing function that pushes the replica to higher interfaces. Selecting the (unknown) crossing probability as the biasing function would ensure *equal* sampling of all interfaces, which is close to optimal. While the crossing probabilities are initially unknown, an iterative procedure can be used to adapt the bias during the simulation, as each interface ensemble naturally gives an estimate for the crossing probability [61, 74]. SRTIS can easily be extended to include multiple states [61] or utilize multiple independent walkers [58, 75].

## III. NOVEL CONCEPTS IN OPS

OpenPathSampling contains many new approaches to implementing transition path sampling simulations, but there are two points that we would particularly like to draw attention to: (1) the use of volume-based interface definitions in TIS and (2) the general treatment of path ensembles.

### A. Volume-based interface definitions

In the original TIS algorithm and most path sampling algorithms based on TIS, interfaces are defined as hypersurfaces in configuration space. To belong to the interface ensemble, a path needs to cross this interface, meaning that at a certain time it is at one side ofthe interface, while a timestep later it is on the other side. We consider a novel interface definition in OPS which relies on hypervolumes in configuration or phase space rather than hypersurfaces. We use the convention that the initial state is inside the hypervolume. In this definition, a path belongs to an interface ensemble defined by a hypervolume if it starts in the initial state, leaves the hypervolume at some point along the path, and terminates in any stable state. The advantage of using volumes instead of surfaces is that set logic (e.g., a union or intersection) can be applied to generate new volume definitions from existing volumes. Fora more extensive discussion see the companion paper [45].

**FIG. 1.**
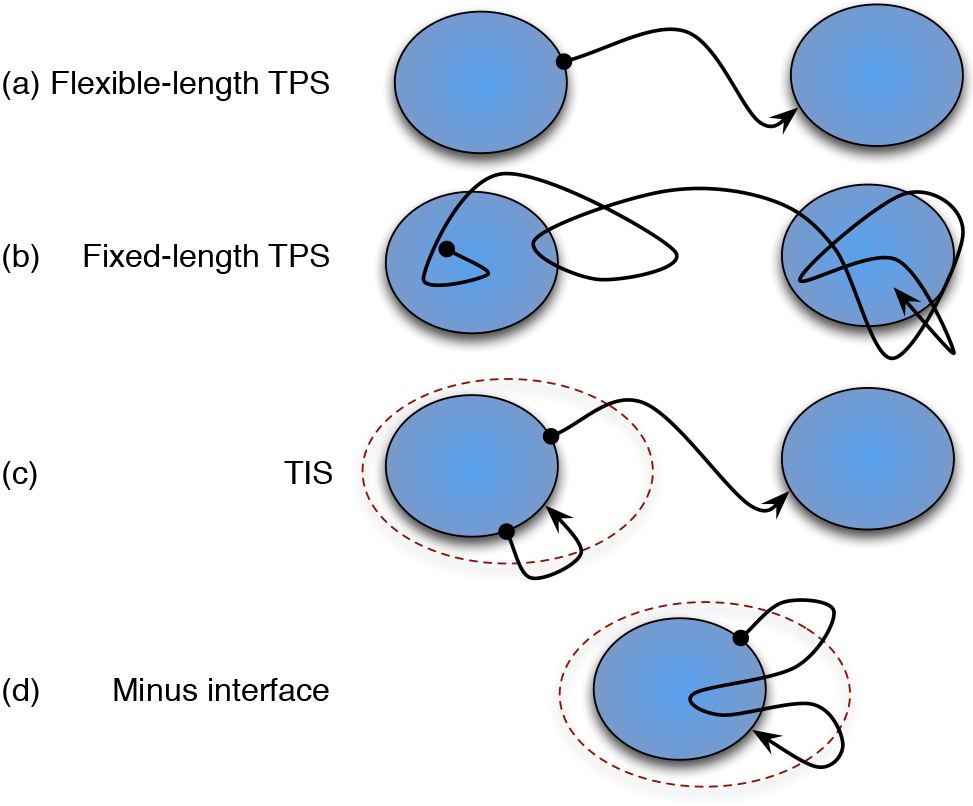
Common path ensembles in TPS and TIS with representative trajectories. Shaded areas represent states, and dashed lines represent interface boundaries.

### B. General treatment of ensembles

One ofthe novel approaches in OPS is the generalization of path ensembles. Previously, each path ensemble had to be treated with specialized code. However, as the number of path ensembles types has grown, the need to treat them in a general fashion arose. In this paper, we make use of a range of path ensembles, including the following, which are illustrated in Fig. 1:

- *Flexible length TPS ensemble* (Fig. 1a): The standard TPS ensemble is a path ensemble between two states. Only the initial and final frames are inside the states.
- *Fixed length TPS ensemble* (Fig. 1b): As with the flexible length TPS ensemble, the initial and final frames must be in the initial and final states. However, the fixed length ensemble has a predefined length, and also allows frames other than the first and final to be in the state.
- *TIS ensemble* (Fig. 1c): The elementary path ensembles in TIS have an interface associated with them. They must begin in a given state, exit the interface hypervolume, and end in any stable state.
- *Minus (interface) ensemble* (Fig. 1d): Paths in the minus ensemble can be described in terms of three segments: the first and last segments are similar to TIS ensemble paths. They start in the state, exit the interface hypervolume, and return to state (where TIS ensemble paths can go to another state, these segments cannot). These two segments are connected by another segment that never exits the interface. Note that this implementation ofthe minus interface ensemble is based on Ref. 71, as opposed to the original minus interface ensemble introduced in Ref. 68. The two versions differ slightly (with the original being subtrajectories of the version used here), however both versions serve the same purpose.

All of these common ensembles can be generalized for more complicated reaction networks. The TPS ensembles become multiple state TPS ensembles if they allow *any* state to be the initial or final state, as long as the initial and final states are different. The TIS ensemble becomes a multiple state TIS ensemble by allowing any state as the final state. The minus ensemble becomes the multiple interface set minus ensemble by taking its interface as the union of innermost interfaces.

OPS allows complicated ensembles to be built from simpler ones. It generalizes both the procedure for testing whether a given trajectory satisfies the ensemble and the procedure for generating new trajectories. Details of this implementation, as well as novel approaches to analysis that this implementation enables, will be discussed in the companion paper [45].

## IV. THE INGREDIENTS OF OPS

Before explaining the OpenPathSampling framework and workflow in more detail, we first explain the frequently used basic objects of OPS that are related to path sampling concepts described in the previous sections. The objects in OPS are divided in two main categories: (1) Data objects that contain the sampled paths and information about the sampling process; and (2) Simulation objects that perform the sampling. All objects generated in OPS, both data and simulation objects, are stored in a single `Storage` file, and can be accessed from it. For example, the `MCStep` objects saved during the simulation can be accessed with `storage.steps` once a file is loaded into `storage`.

### A. Data objects

The main data objects of OPS fit into a hierarchy as shown in Fig. 2. The data structure can be divided into *what* is being sampled (i.e., which trajectories from which ensembles), and *how* it is being sampled (i.e., the nature of the path moves performed.) All of this is unified in the MCStep object, which describes a step of the path sampling simulation, and which has two important attributes: a `SampleSet` object called `active`, which records the state of all replicas in the simulation at the end of a given simulation step (the “what”); and a `MoveChange` object called change, which describes what happened during the simulation step (the “how”). Below we describe these attributes in more detail.

**FIG. 2.**
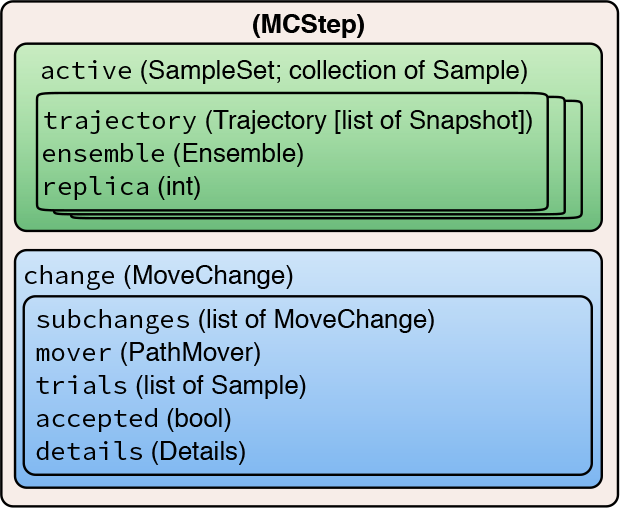
Hierarchical data structure of the MCStep data object. The attribute names are shown, and the type is provided in parentheses.

#### 1. Data structures for what is being sampled

- `Snapshots`, sometimes called “frames” or “time slices,” are at the core of any simulation technique. They describe the state of the physical system at a point in time, and in molecular dynamics, typically consist of coordinates, velocities, and periodic cell vectors. The Snapshot object in OPS can be easily extended to carry additional data, such as wavefunction information or variables from an extended phase space.
- A `Trajectory`, also called a “path,” is essentially a list of `Snapshots` in temporal order. In addition, it provides several convenience methods, for example, to identify which `Snapshots` are shared by two trajectories.
- The `Sample` object is a data structure that links a `Trajectory` with the Ensemble object (described in section IV B) from which it was sampled, and an integer replica ID. The `Sample` is needed because methods such asTIS, and especially RETIS, sample multiple ensembles simultaneously. Correct analysis requires knowing the ensemble from which the `Trajectory` was sampled.
- Since methods likeTIS have several active `Samples` during a path simulation step, OPS collects them into one SampleSet. The `SampleSet` contains a list of `Samples`, and also has convenience methods to access a sample either by replica ID or by ensemble, using the same syntax as a Python `dict`.

#### 2. Data structures for how the sampling occurs

- The `MoveChange` contains a record of what happened during the simulation step. Because the simulation move itself generally consists of several nested decisions (type of move, which ensemble to sample, etc.),the `MoveChange` object can contain subchanges, which record this entire sequence of decisions. In addition, it includes a pointer to its `PathMover` (described in section IV B), a list of the trial `Samples` generated during the step, and a boolean as to whether the trial move was accepted.
- The `MoveChange` also contains a `Details` object, which is essentially a dictionary to store additional metadata about a move. This metadata will vary depending on the type of move. For example, with a shooting move, it would include the shooting point. In principle, all the additional information that might be of interestforanalysisshould be stored in the `Details`.

**FIG. 3.**
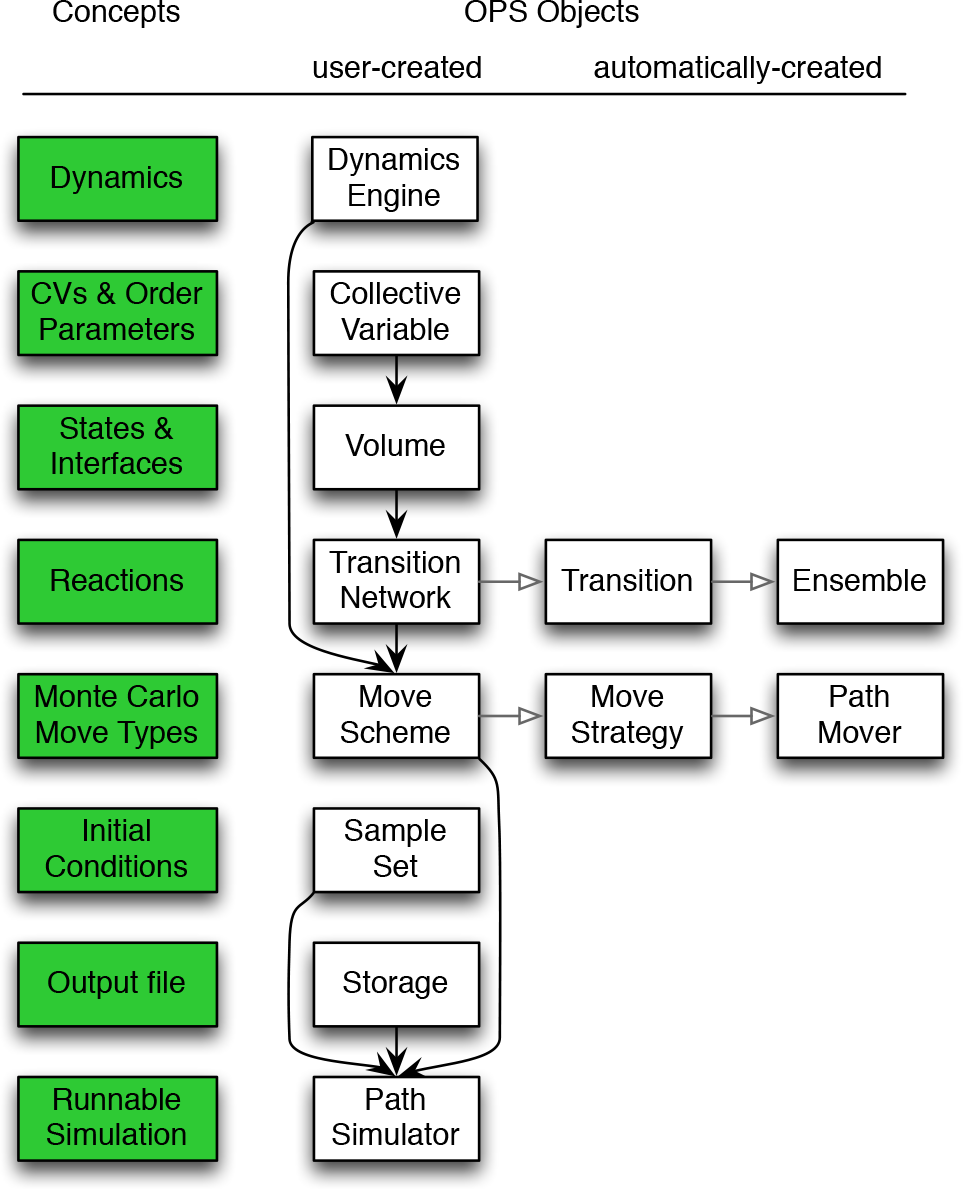
Schematic representation of the connection between the path sampling concepts and their related OPS objects. The concepts are listed in the leftmost column, shaded green. The next column shows the objects which must be created by a user to run a simulation. The filled arrows indicate when one object is the input to create another object. The objects in the right two columns are automatically created. The open arrows point from an object to the objects it automatically creates. In this way a `TransitionNetwork` creates `Transition` object that creates in turn `Ensemble` objects.

### B. Simulation objects

The simulation objects actually perform the simulation, and can be assembled in different ways to perform many types of simulations. In addition, simulation objects in OPS can be stored. This facilitates restarts to continue a simulation and enables re-use for other types of simulations, e.g., using the same state definitions for committor analysis as well as path sampling. The `PathSimulator` class contains all the information to run the simulation. The `PathSampling` subclass of `PathSimulator` is used for path sampling simulations. Fig. 3 shows the relation between path sampling concepts and the associated objects in OPS. Each of the components is described in more detail below.

- A `DynamicsEngine` performs the actual molecular dynamics: that is, it generates a trajectory from an initial frame. OPS has built-in support foran internal toy dynamics engine (primarily intended for2D models) and for OpenMM [76]. Support for Gromacs [77, 78] and LAMMPS [79] will be added in future releases.
- A `CollectiveVariable` is a function of a Snapshot, and in many cases is just a function of the coordinates. It is also sometimes called the “order parameter,” “progress variable,” “reaction coordinate,” or “feature.” In line with the rare event terminology (e.g., [80, 81]) the neutral term CV (for Collective Variable) can both be used to define interfaces and states (via `Volumes`), as well as to construct order parameters. The `CollectiveVariable` in OPS is a wrapper class around an arbitrary function. For example, the `CoordinateFunctionCV` will wrap any user-defined function that only depends on the snapshot’s coordinates. In addition, specific classes enable the use of functions from other packages, e.g, the `MDTrajFunctionCV` providesa wrapper class forfunc-tionfrom the MDTraj[82] analysis package. Otherwrappers exist for MSMBuilder [83, 84] and PyEMMA [85].
- The `Volume` class in OPS represents a hypervolume in phase space. This can be used to define a *state*, also called a “core set.” In addition, *interfaces* are also defined by volumes, rather than by hypersurfaces as in the traditional TIS literature (see section III A). A volume is typically defined based on allowed ranges of CVs; in OPS the `CVDefinedVolume` object creates such a volume based on a minimum and maximum value of the CV.
- The `Ensemble` class in OPS defines the paths that are allowed within a given path ensemble. It is more accurately thought of as the indicator function fora restricted path ensemble (c.f. Eq. 3). The indicator function alone reduces the set of all possible paths to the trajectories with non-zero probablity in the path ensemble, but with no distinction in their relative statistical probabilities. Sampling according to the correct statistical weights is the role of the `PathMover`, described below.

In addition to the indicatorfunction, `Ensemble` objects contain two methods, `can_append` and `can_prepend`, which check whether a given trajectory could be appended or prepended into a trajectory in the ensemble. This allows us to create a rich toolkit to create custom ensembles. For instance, a path that connects states *A* and *B* is defined as a trajectory that follows the sequence ofeventsthatit is first in *A*, then notin (*A* ∪ *B*), and finally in B. In OPS, this sequence is described with a `SequentialEnsemble` object, which provides a flexible way to implement arbitrarily complex path ensembles (see Ref. [45]).

Despite this powerful toolkit and the fundamental role of the `Ensemble`, under most circumstances the user does not need to instantiate `Ensemble` objects. Instead, they are automatically created by the `Transition` and `Network` objects, described below.

- A `Transition` object contains all information for studying a single-direction reaction connecting a specific initial state and a specific final state, such as *A* → *B*, and serves as an organizational structure for systems with many states, where the numberof possible transitions grows as *N*(*N*-1) forNstates. ForTPS, this object consists just of one ensemble, while forTIS it usually consists of several interface path ensembles, as well as the minus ensemble (used in RETIS). Note that *A* → *B* and *B* → *A* are two different transitions, each with their own sets of ensembles, thus requiring two `Transition` objects. A single rate *k* would be associated with each `Transition` and *k*_*A*→*B*_ ≠ *k*_*B*→*A*_.
- A `TransitionNetwork` object (which we will frequently refer to as simply the “network”) consists of a set of `Transitions`. Since OPS is designed to handle the systems with many states, the network gathers all the transitions into one object. It is a network in the graph theory sense: states are nodes; reactions (transitions) are directed edges. Subclasses of `TransitionNetwork`, such as `TPSNetwork` or `MSTISNetwork`, deal with specific approaches to sample the network. All the ensembles to be sampled are contained in the `TransitionNetwork`. Section VD provides more details.
- `PathMovers`, or“movers,” perform Monte Carlo moves in path space, such as shooting, reversal, minus, or replica exchange. They are organized into a *move decision tree*, which selects the specific move to use (the move type and the ensemble). An example of a move decision tree is given in Fig. 6. The `Ensemble` associated with a given mover determines whether a trajectory is in the path harvest forthat mover, butthe mover itself can reject paths such that the correct statistics forthe path ensemble are obeyed (i.e., to preserve detailed balance.) `PathMovers` are discussed in more detail in section V E1.
- The `MoveScheme` contains and builds the move decision tree, which in turn contains all the `PathMovers` available to a simulation. The `MoveScheme` is created by associating several `MoveStrategy` objects with it. Each `MoveStrategy` builds several related `PathMovers`. For example, a `NearestNeighborReplicaExchangeStrategy` will create a `ReplicaExchangeMover` for each pair of nearest-neighbor ensembles in each Transition from the TransitionNetwork. Options for creating the strategy can control which ensembles are used, and whether this adds to or replaces existing strategies. This provides the user a great deal of flexibility when customizing the move decision tree using the `MoveScheme` and `MoveStrategy` objects. For simplicity, OPS provides a `DefaultScheme` with reasonable defaults forTIS (one-way shooting, nearest-neighbor replica exchange, path reversal, and minus move), and a `OneWayShootingMoveScheme` with a reasonable default for TPS. The `MoveScheme` and `MoveStrategy` objects will be discussed in more detail in section V E 2.

**FIG. 4.**
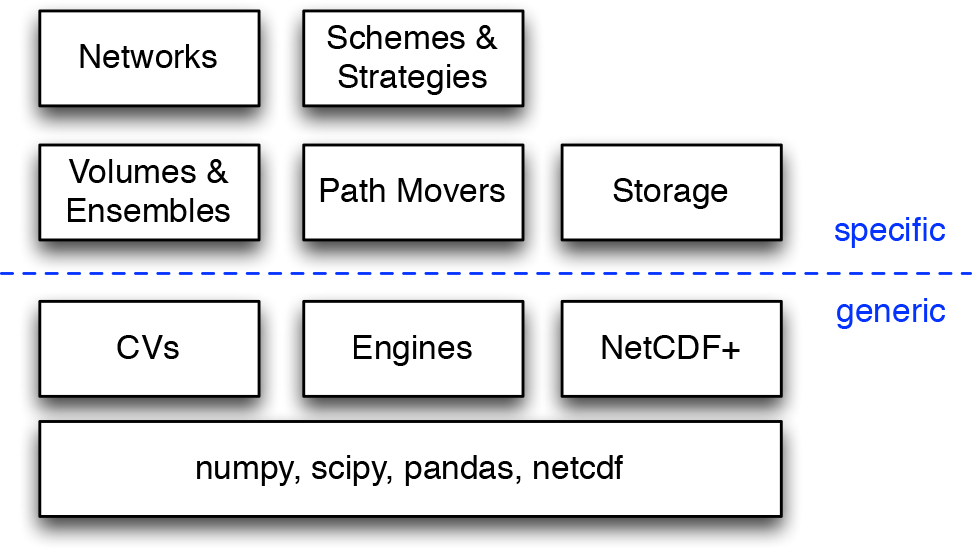
The modules of OPS can be separated into different layers of abstraction. The layers can be considered as both increasing specificity of purpose (from bottom to top) as well as increasing ease of use or ease of implementation of new subclasses. Underneath the OPS modules the are external packages upon which OPS is built. Above that are OPS modules which have potential for use outside the context of reaction dynamics and path sampling. Above that the code becomes more specific to path sampling, and to the OpenPathSampling project. At the top layer, some ofthe more powerful OPS libraries are abstracted into a more simple user interface. The level of user that is likely to spend significant time working at each level is indicated on the left.

### C. Layers of abstraction in OPS

OPS is structured as a set of Python modules, organized according to major classes. As a library, users can interact with different levels of abstraction. Fig. 3 and the previous section havealready indicated how `TransitionNetworkobjects` act as a more user-friendly layer for `Ensembles`, and how `MoveScheme` and `MoveStrategy` objects create a simpler layer for working with `PathMovers`. But these lower-level objects can also be accessed by users, as will be discussed in the companion paper [45].

Objects like `Ensembles` and `PathMovers` are specific to path sampling and related topics. These are built on even more generic objects, which might be useful beyond the scope of path sampling. Many `PathMovers` use the generic `DynamicsEngine` wrapper to run the molecular dynamics. Volumes are defined in terms of `CollectiveVariables`, which have many uses beyond path sampling. The specific OPS `Storage` class is based on more generic `NetCDFPlus` subpackage, built for OPS. This is shown in Fig. 4, where lower levels are more generic, while higher levels are more specific to path sampling. Higher levels also tend to be more user friendly.

## V. OPS WORKFLOW

In this section we give an overview ofthe process for setting up and running a path sampling simulation with OpenPathSampling, including some general discussion on practical aspects of path sampling simulations. In general, every path sampling simulation can be split into the following steps:

1. Settingupthe moleculardynamicsengine
2. Defining states and interfaces
3. Setting up the transition network and move scheme
4. Obtaining initial pathways
5. Equilibration and running the simulation
6. Analyzing the results

In practice, the human effort in path sampling using OPS will focus on defining the states and interfaces, obtaining trajectories for initial conditions, and analyzing the simulation results. OPS aims to facilitate those steps and automate what it can, such as setting up the `TransitionNetworks` and `MoveSchemes`, and running the simulation. In addition, OPS provides many tools for the analysis ofthe simulation results.

In the first setup steps (1-3), the user chooses the dynamics of interest and decides on the `DynamicsEngine`, the `CollectiveVariables`, defines the `Volumes` for the states and interfaces, as well as the topology of the reaction network, and decides on the sampling `MoveScheme`. Fig 3 renders these steps from the top down. Selecting relevant collective variables and using them to define state volumes is of critical importance, but is also dependent on the system being studied. We assume that a user is already familiar enough with the system to make reasonable choices for these.

The specific definition ofthe transition network is handled in OPS by a `TransitionNetwork` object, which automates the creation of `Transitions` and `Ensembles` for common variants of TPS (including multiple state) and TIS (including multiple state and multiple interface set variants). These objects take as input the `Volume` based states and interfaces definitions.

The `MoveScheme` is created based on the `TransitionNetwork` and a `DynamicsEngine`. It can be customized by adding additional `MoveStrategy` objects, but OPS provides default schemes for convenience. The MoveScheme and its accompanying `MoveStrategy` objects create all the PathMovers. Each `PathMover` knows on which ensemble(s) it acts, and are organized into a total move decision tree.

The final initialization step is to create an initial `SampleSet` by loading valid preexisting initial trajectories into each of the ensembles. See Appendix A for several approaches to obtain initial conditions.

The simulation is performed by a `PathSimulator` object. Path sampling simulations use a subclass called `PathSampling`. Other subclasses of the `PathSimulator` include `CommittorSimulation` for calculating committors and `DirectSimulation` for calculating rates and fluxes via direct MD. All `PathSimulator` objects take a `Storage` object as input, to determine where to save data. In addition, `PathSampling` takes the `MoveScheme` and the initial `SampleSet` as input.

Analysis is done independently from the sampling and requires only the `Storage` and `TransitionNetwork` for the computing observables, and additionally the `MoveScheme` for the sampling statistics. Everything that is needed for analysis is stored in the output file, including the `TransitionNetwork` and `MoveScheme`.

In the next subsections we discuss these six steps in more detail.

### A. Step 1: Setting up the molecular dynamics

Of course, before embarking on a path sampling simulation, one must decide on the system to simulate, and the nature of the underlying dynamics (i.e., the thermodynamic ensemble represented, the integrator used for the dynamics, the force field to define interactions, etc.) OPS is designed to wrap around other engines to take advantage of the flexibility already built into other software. Currently, OPS supports OpenMM [76] as well as its own internal dynamics engine intended mostly for2D toy models.

The basic `Engine` takes general OPS specific options defining, e.g., handling of failing simulations, maximal trajectory length, etc, as well as dimensions used in snapshots that the engine generates (e.g., number of atoms). Each specific engine also carries information necessary for it to setup a simulation. In case of OpenMM this includes a description of the `Integrator`, the `System` object (force field, etc.), the systems `Topology` and some OpenMM specific options (e.g., hardware platform and numerical precision).

### B. Step 2: Defining states and interfaces

The ensembles used in path sampling methods require definitions of (meta)stable states and, in the case of transition interface sampling, interfaces connecting these states. OPS implements both states and interfaces in terms of `Volume` objects.

The main types of volume objects are the `CVDefinedVolume`, and its periodic version, `PeriodicCVDefinedVolume`. Each of these defines a volume in phase space based on some `CollectiveVariable`. This could include such quantities as atom-atom distances, dihedral angles, RMSD from a given reference frame, number of contacts, etc. The user must first define a `CollectiveVariable` object, either as a wrapper around functions from other software packages (some examples below use `MDTrajFunctionCV`, which wraps MDTrajanalysis function), oraround a user-written function (other examples will show the use of the more general `CoordinateFunctionCV`).

Using the `CollectiveVariable` we have a clear seperation between the full simulation data and what we consider relevant for state definitions and later analysis. Thisseperation allows us to later run analysis without the need to load a single frame, and to store a reduced set without the actual coordinates.

To define a volume, the user must also specify minimum and maximum values for the CV. The volumes can then be created with, e.g., `CVDefinedVolume(cv, minimum, maximum)`, which defines a `frame` as being inside the volume if `minimum ≤ cv(frame) ≤ maximum`. Volumes can be combined using the same set operation as Python sets: &(intersection), | (union), - (relative complement), ˆ (symmetric difference), and ~ (complement). Volume combinations of the same collective variable are automatically simplified when they can be recognized [e.g., (0 ≤ *x* < 5)&(3 ≤ *x* < 8) becomes 3 ≤ *x* < 5]. The ability to arbitrarily combine volumes allows one to define arbitrary states, e.g., “this hydrogen bond is formed and this dihedral is near a certain value.” This provides OPS with powerful flexibility.

### C. Step 3: Setting up the transition network and move scheme

The transition network (path ensembles) and the move scheme (Monte Carlo moves) can be thought of as *what* to sample, and *how* to sample, respectively.

For complex TIS simulations, the number of path ensembles to be sampled can grow into the hundreds. `TransitionNetwork` objects efficiently create those ensembles according to standard ways of organizing, and facilitate later analysis. The examples in Sec. VI will demonstrate the four main kinds of network objects: `TPSNetwork` for flexible-length TPS, `FixedLengthTPSNetwork` for fixed-length TPS, `MSTISNetwork` for multiple-state TIS, and `MISTISNetwork` for TIS and multiple interface set TIS.

The `MoveScheme` createsand organizesthe possible Monte Carlo moves, as appropriate for a given transition network. As with the transition networks, the `MoveScheme` object also facilitates later analysis. The examples in section VI will go over the simplest default move schemes (`OneWayShootingMoveScheme` for TPS; `DefaultScheme` for TIS). However, the move scheme is very customizable, as will be elaborated on in the companion paper [45].

As both the `TransitionNetwork` and `MoveScheme` are crucial in OPS, we devote extra attention to these objects below.

### D. Step 3a: Transition networks

The `TransitionNetwork` object contains all the path ensembles to be sampled for the reaction network of interest. To simplify analysis, most ensembles are grouped into `Transition` objects, which describe a single transition within the network. There are also special ensembles (e.g., ensembles associated with multiple state interfaces or with minus interfaces) which may not be specific to a single transition, and are only associated with the network as a whole. In general, the user only needs to create the `TransitionNetwork` object, which will automatically create the relevant `Transitions` and `Ensembles`. The simplest transition network contains a single transition, the one-way *A* → *B*. A bidirectional network *A* ↔ *B* is thus characterized by two transitions, each associated with its own set of ensembles.

**TABLE I.** Predefined network types and the number of (physical) transitions, sampling transitions, and sampling ensembles arising from different initialization parameters. Volumes are represented with capital letters (e.g., `A` or `B`) and interface sets are represented as `mA` for the interfaces leaving `A` in MSTIS, or `mAB` for interfaces leaving `A` toward `B` in MISTIS. The number of interfaces in an interface set is given by *m*_*A*_ or *m*_*AB*_, respectively. The total number of states is assumed to be *N* (with the final state represented by `N`).

| Network and Initialization | Transitions | Sampling Transitions | Sampling Ensembles |
| --- | --- | --- | --- |
| **TPSNetwork:** |  |  |  |
| A, B | 1 | 1 | 1 |
| [A, B], [B, C] | 3 | 1 | 1 |
| [A, B, ..., N], [A, B, ..., N] | $N(N-1)$ | 1 | 1 |
| **MSTISNetwork:** |  |  |  |
| [(A, mA), (B, mB)] | 2 | 2 | $m_A + m_B$ |
| [(A, mA), (B, mB), ..., (N, mN)] | $N(N-1)$ | $N$ | $\sum_I m_I$ |
| **MISTISNetwork:** |  |  |  |
| [(A, mAB, B)] | 1 | 1 | $m_{AB}$ |
| [(A, mAB, B) (A, mAC, C), ..., (A, mAN, N)] | $N-1$ | $N-1$ | $\sum_{J \neq A} m_{AJ}$ |
| [(A, mAB, B), ..., (A, mAN, N), ...<br>..., (N, mNA, A)] | $N(N-1)$ | $N(N-1)$ | $\sum_I \sum_{J \neq I} m_{IJ}$ |

Each network involves two groupings of transitions: the *sampling transitions* and the *physical transitions*. MSTIS shows a clear example of the distinction between these: while sampling, the transitions studied are *A* → (*B* ∪ *C*), *B* → (*A* ∪ *C*), and *C* → (*A* ∪ *B*). However, in analysis we obtain the rates for all the individual physical transitions *A* → *B*, *A* → *C*, *B* → *A*, *B* → *C*, *C* → *A*, and *C* → *B*. For a network with *N* states, up to *N*(*N* - 1) unique physical transitions are possible. The sampling transitions are found in a `list`, accessed as `network.sampling_transitions`, and the physical transitions are in a `dict`, with state pairs (`initial, final`) as keys, and the associated `Transition` object as value.

The `Transition` and the `TransitionNetwork` objects depend on the type of simulation that is intended, just as the `Ensemble` does. Table I shows how different input parameters create different numbers of physical and sampling transitions for the built-in network objects in OPS. The network fora TPS simulation is made with either the `TPSNetwork` or `FixedLengthTPSNetwork` objects. The `TPSNetwork` is initialized with a list of initial states and a list of final states; all pairs of (non-self) transitions are generated internally. A `TPSNetwork` has only one sampling `TPSTransition`, which has only one ensemble. However, for analysis the network includes ensembles for every possible physical transition. If `A` is the only initial state and `B` is the only final state, then *A* → *B* is the only physical transition. When multiple initial and final states are given, then all the non-self physical transitions are allowed: in the second line of Table I, that would be *A* → *B*, *A* → *C*, and *B* → *C*. When all *N* states are given as both initial and final states, all *N*(*N* - 1) non-self transitions are included. The `FixedLengthTPSNetwork` is exactly like the `TPSNetwork`, except that its initialization also requires the length of the path (in snapshots).

Within standard TPS approaches, there is a one-to-one correspondence of (sampling) ensemble to network. That makes these networks relatively simple. The situation becomes more complicated with TIS. In TIS, each transition involves a set of interface ensembles. In addition, there are the minus ensembles, which (in MISTIS) can be associated with more than one transition, and there are the multiple state outer ensembles (in MSTIS and MISTIS), which are also associated with more than one transition.

The `MSTISNetwork` and `MISTISNetwork` are initialized with specific data about the transitions. In MSTIS, this includes the initial states and the interface sets associated with them, provided as a `list of tuples`. The `MSTISNetwork` creates sampling ensembles that allow paths that end in any state, and always samples all transitions between all states. As shown in Table I, it therefore always has *N*(*N* - 1) physical transitions and *N* sampling transitions for *N* input states. The number of ensembles depends on the number of ensembles per interface set, but scales linearly with the number of states.

In addition to the initial states and the interface sets, `MISTISNetwork` also requires the ending state for each transition, provided asa third item in each tuple. The numberof physical transitions for the `MISTISNetwork` is always equal to the number of sampling transitions, and the number of ensembles grows with the number of sampling transitions. This means that, in the worst case of sampling all possible transitions, the number of ensembles scales quadratically with the numberof states. However, MISTIS has the advantage that it allows one to select only specific transitions of interest, or to use different interface sets for transitions beginning in the same initial state, allowing each transition to be sampled more efficiently.

Both of these TIS networks automatically create appropriate minus interface ensembles, and they can optionally take an `MSOuterTISInterface` for the multiple state (MS) outer interface ensemble. The MS-outer ensemble is the union of several TIS ensembles starting from different initial states [34, 71]. Whereas a TIS ensemble only allows trajectories that begin in a single given initial state, the MS-outer ensemble allows trajectories that begin in any of multiple initial states. This ensemble, combined with replica exchange, facilitates decorrelation of trajectories.

MSTIS and MISTIS are two different ways to create ensembles to study a reaction network. The MSTIS approach is more efficient when all transitions from the same state are described by the same order parameter. The MISTIS approach allows more flexibility in sampling, by allowing different transitions from an initial state to use different order parameters or selection of specific transitions of interest.

The simplest network, *A* → *B*, can be studied using the `MISTISNetwork` object. The bidirectional *A* ↔ *B* network can be studied using either a `MISTISNetwork` or a `MSTISNetwork`: the ensembles which are created would be indistinguishable.

These networks are not exhaustive, and other possibilities might be implemented by users. For example, it might be interesting to sample transitions from one state to all other states in an MSTIS simulation. This cannot be done with the built-in `MSTISNetwork`, but it would be relatively straightforward to create another subclass of `TransitionNetwork` that allows this.

### E. Step 3b: The Monte Carlo Move Scheme

#### 1. Path movers

In OPS, each `PathMover` instance is connected to specific ensembles. For example, there is a separate shooting mover for each ensemble, and a separate replica exchange mover for each pair of ensembles that are allowed to swap in replica exchange. The `move` method ofthe `PathMover` object actually performs the Monte Carlo move. It takes a `SampleSet` as input, and returns a `MoveChange`, which the `PathSimulator` applies to the original `SampleSet` in order to create the updated `SampleSet`.

OPS includes a rich toolkit so that developers of new methods can create custom methods. Those toolkits are discussed in detail in the companion paper [45]. Here, we will introduce some ofthe built-in path movers.

- *Shooting movers*: OPS has support for both oneway (stochastic) shooting [48, 65] as well as the twoway shooting algorithm. These are implemented as `OneWayShootingMover` and `TwoWayShootingMover`. In addition to a specific ensemble, the shooting movers require a `ShootingPointSelector` to choose the shooting point. The most commonly used selector is initial the `UniformSelector`, which selects any point except the endpoints of the trajectory (which are in the defined states) with equal probability. Other possibilities could also be implemented, such as using a Gaussian distribution [23] or a distribution constrained to the interface [69]. The `TwoWayShootingMover` also requires a `SnapshotModifier` to change the snapshot in some way (e.g., modifying the velocities). Several possibilities exist, includingeitherchangingthe direction ofthe velocity for some atoms, or completely randomizing velocities according to the Boltzmann distribution.
- *Path reversal mover*: Another standard mover is the `PathReversal` mover, which takes the current path in the ensemble and tries to reverse its time direction. For a path that leaves and returns to the same stable state this move is always accepted. As stated in Sec. IIC, this move helps to decorrelate the sampled trajectories.
- *Replica Exchange mover*: A `ReplicaExchangeMover` involves two ensembles (see Fig. 6). When a move is attempted, the mover takes the paths associated with these ensembles in the current sample set, and tries to exchange them. This trial move will be accepted if both paths are valid paths in their respective ensembles.
- *Minus mover*: The `MinusMover` is a more complicated `PathMover`. In essence, it combines replica exchange with extension of the trajectory. OPS has a toolkit to simplify the creation of more complicated moves from simpler ones, which is be discussed in more detail in the companion paper. The `MinusMover` uses both the minus interface ensemble and the innermost normal TIS ensemble. It extends the trajectory from the innermost ensemble until it again recrosses the interface and returns to the stable state, resulting in a trajectory with two subtrajectories that satisfy the innermost TIS ensemble. This trajectory satisfies the minus ensemble. The trajectory that had previously been associated with the minus ensemble also has two subtrajectories that satisfy the innermost TIS ensemble, and one of them is selected. After the move, the newly extended trajectory is associated with the minus ensemble, and the selected subtrajectory is associated with the innermost TIS ensemble.

**FIG. 5.**
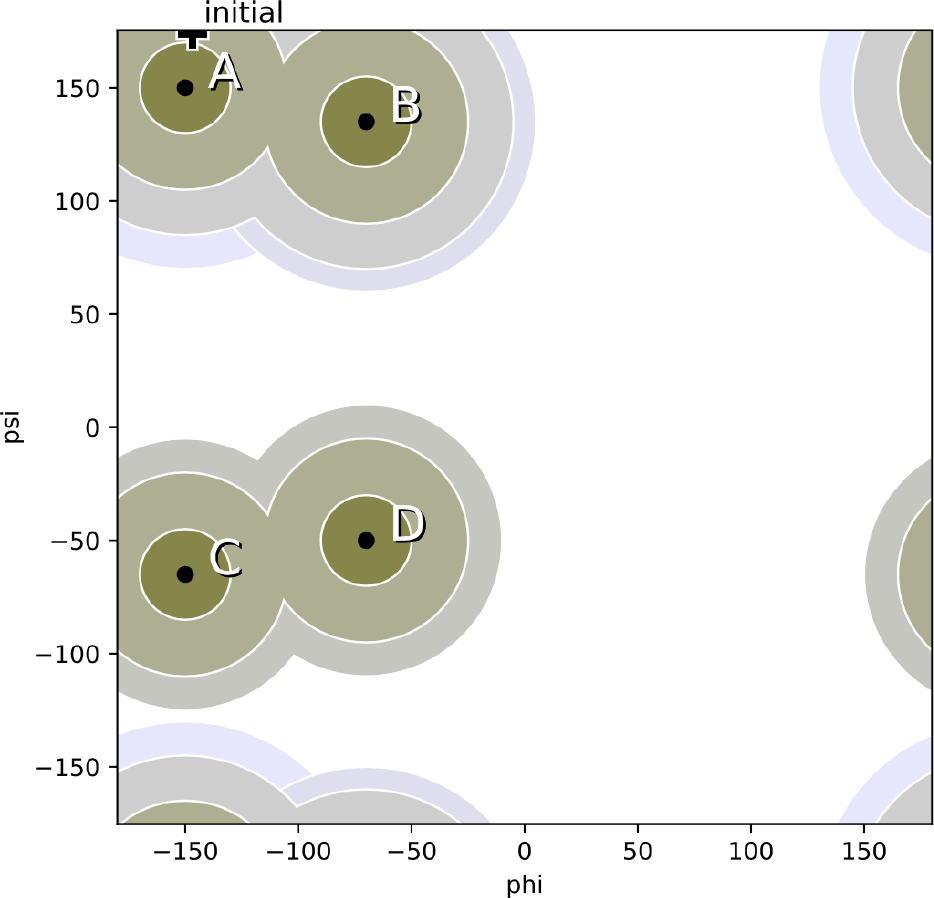
Example of a transition network used in the MSTIS alanine dipeptide examples (See Section VI) Multiple states A-D are defined according to the dihedral angles *ψ* and *ϕ*. THe core sets for A-D are defined as being within 10 degrees of the core center (indicated by black dot). Each state has its own set of interfaces using the geometric distance in *ψ* - *ϕ* space to the core center, indicated by shadede circles. The `MSTISNetwork` object creates for each state the collection of path ensembles for each interface, plus the minus interface. In addition there is a multiple state union interface for the outermost interfaces. The plus marks the location of the initial conformation used in the example.

#### 2. The MoveScheme and MoveStrategy

The `MoveScheme` creates and contains the move decision tree, which is essentially the protocol for the simulation. Fig. 6 shows a graphical representation of the decision tree created by a simple `MoveScheme`. The decision tree contains the different choices of move type (e.g. shooting, reversal, replica exchange) and assigns specified weights to them. At the leaves of the tree are path movers. Each path mover acts on a certain ensemble (shown on the right of Fig. 6).

The `MoveScheme` object organizes the path movers in several *mover groups*, held in a dictionary called movers, with strings as keys and a list of `PathMovers` are values. Each group corresponds to a related set of movers (which are used on different ensembles). For example, the default shooting movers are in the group ‘`shooting`’ and thedefault replica exchange movers are in the group ‘`repex`’.

The most common `MoveScheme` objects are the `DefaultScheme` (for TIS) and the `OneWayShootingMoveScheme` (for TPS). All move schemes require a network; `DefaultScheme` and `OneWayShootingMoveScheme` also require an engine. The move decision tree can also be generated by hand, and then given as input to a `LockedMoveScheme`, although some additional information (such as the `choice_probability`, a dictionary mapping each path mover to its relative probability of being selected) must be manually added to a `LockedMoveScheme` for some analysis to work. Furthermore, a `LockedMoveScheme` cannot by modified using `MoveStrategy` objects.

In general, the easiest way to customize the move scheme is to start with a `DefaultScheme` or a `OneWayShootingMoveScheme`, and then append strategies that give the desired behavior. The whole scheme is built by applying the `MoveStrategy` objects in sequence. Each subclass of `MoveStrategy` has a priority level associated with it, and the strategies are built in an order sorted first by that priority level, and second by the order in which they were appended to the scheme (so later additions can override earlier versions). Several aspects of the way a `MoveStrategy` contributes to the move decision tree can be set in its initialization: which ensembles the strategy applies to, which mover group the strategy is for, and whether to replace the effects of previous strategies. Additionally, mover-specific parameters (such as shooting point selector forshooting moves) are passed along to the movers that are constructed by the strategy.

**FIG. 6.**
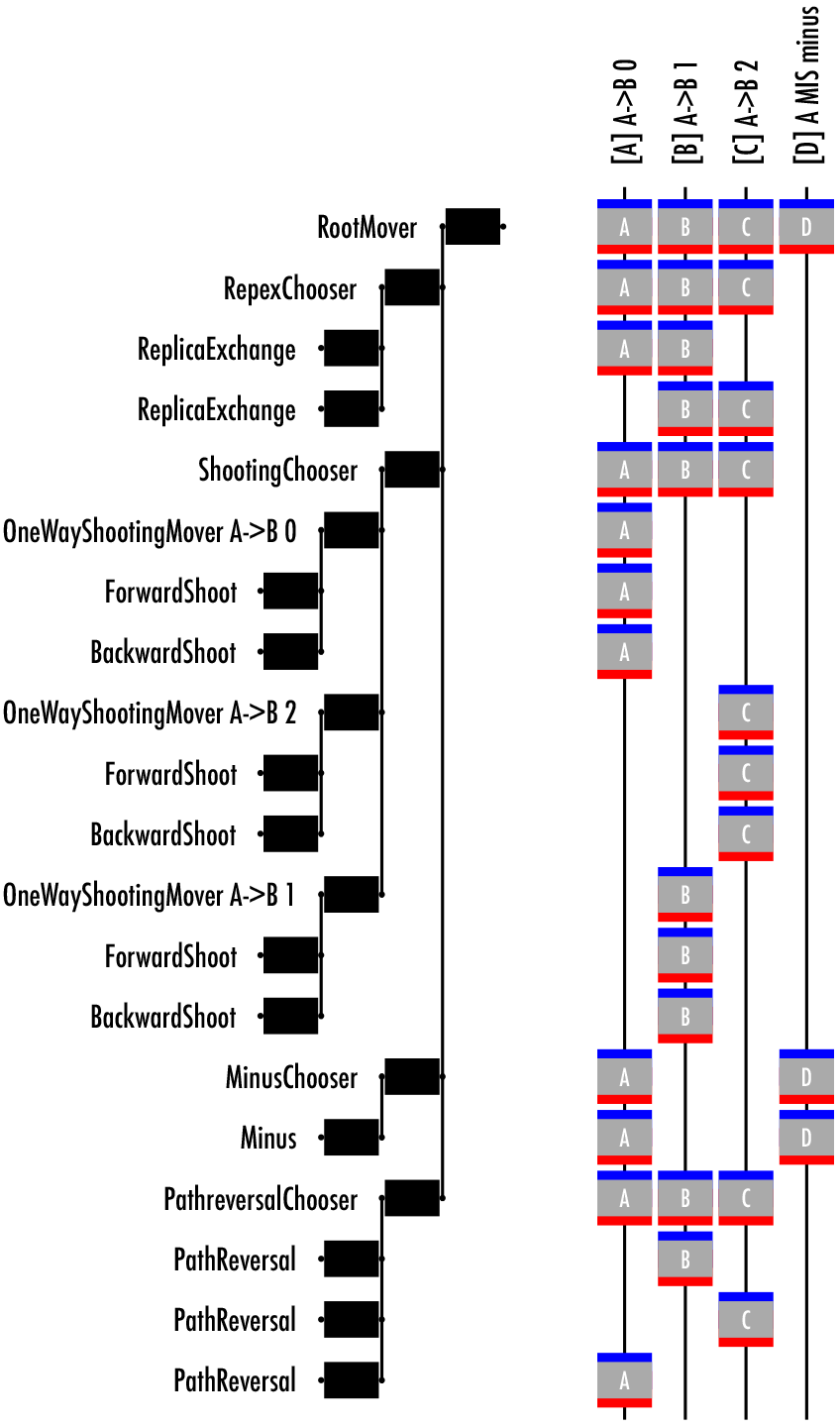
Schematic representation of the decision tree as constructed by the MoveScheme object. Shown is an example for RETIS. The `MoveScheme` points to the root of this tree (left). The branches are the different move levels. First level is the decision about what type of move: shooting, replica, reversal. Next level is the decision about what ensemble needs to be moved. For the shooting, the next level is about which direction the shot is. For other moves the choice is slightly different. The right part of the picture show which ensembles are affected. Each vertical line denotes an ensemble. At the root of the tree each ensemble can be chosen. Going down the tree, the ensembles affected reduce in number. The letters are arbitrary labels for each ensemble. The grey box around each letter show the input (red) and the output (blue). This sort of schematic can be generated using the `paths.visualize.MoveTreeBuilder` object.

This allows one to, for example, add a shooting move of a differenttype (e.g., two-way instead of one-way, or using a different shooting point selection algorithm) fora specific ensemble — either overriding the original mover or adding a second “group” of shooting movers (with a different name, e.g., `‘shooting2’` so as not to conflict with the existing `’shooting’`). One might do this so that there are two kinds of shooting moves: one which causes large decorrelations in path space (but might have lower acceptance) and one that has a better acceptance probability.

### F. Step 4: Obtaining initial conditions

The initial conditions fora path sampling simulation consist of a `SampleSet` with at least one `Sample` for any possible initial move. As discussed above, each `Sample` consists of a trajectory, the ensemble for that trajectory, and a replica ID. Preparing the initial sample set thus breaks down into two parts: (1) creating the initial trajectories; (2) assigning them to appropriate ensembles and giving them individual replica IDs.

The first part, obtaining initial trajectory from the path ensemble, is in general non-trivial since the events we are interested in are rare, and the best approach is likely to be system specific. We discuss several possible approaches in detail in Appendix A.

The second partis much easier. Once we have trajectories, `scheme.initial_conditions_from_trajectories` will take those trajectories and create appropriate initial conditions for the move scheme called `scheme`. This method attempts to create a sample for every ensemble required by the move scheme by checking if the given trajectories (orsubtrajectories of them) or their time-reversed versions satisfy the ensemble. Internally, this uses the ability of the `Ensemble` object to test whether a trajectory (or subtrajectory thereof) satisfies the ensemble.

For some ensembles, such as the minus interface ensemble, the method `extend_sample_from_trajectories` has been implemented, which runs dynamics to create a trajectory that satisfies the ensemble, starting from input subtrajectories.

Last, the move scheme method `MoveScheme.assert_initial_conditions` can be used to check if a given set of initial conditions contains all `Samples` needed to run the simulation, and raises an `AssertionError` if not.

### G. Step 5: Equilibration and running the simulation

As with other simulation techniques, such as molecular dynamics and configurational Monte Carlo, the equilibration process for path sampling is often just a shorter version of the production run. Both equilibration and production require creating a `PathSimulator` object, which create the runnable simulation. The examples here focus on `PathSampling`, but other subclasses of `PathSimulator` include `CommittorSimulation` and `DirectSimulation`(for rates and fluxes). The `PathSampling` simulator is initialized with a storage file, a move scheme, and initial conditions. It has a `run` method which takes the number of MC trial steps to run. All the simulation and storage to disk is done automatically.

### H. Step 6: Analyzing the results

OPS has many built-in analysis tools, and users could create a wide variety of custom analyses: the companion paper includes several examples [45]. However, nearly all analysis of path sampling falls into two categories: either the analysis provides information about the ensemble that is sampled (often tied to observables such as the rate) or the analysis provides information about the sampling process itself. Both analysis types are extremely important — poor behavior of the sampling process would indicate low confidence in the calculated observable. And, of course, combining insights from both can yield understanding of the physical process understudy. The basic use of OPS analysis tools to calculate rates from MSTIS and MISTIS simulations and mechanistic information (path densities) from TPS simulations, as well as properties of the sampling process such as the replica history tree (a generalization of the “TPS move tree” in existing literature), measures of mover acceptance ratios, and measures of the replica exchange network and its efficiency, will be illustrated in the following examples.

## VI. ILLUSTRATIVE EXAMPLES

In this section, we give and discuss several examples. These examples are meant to show the user how to set up, run, and analyze several basic applications of *TPS*, *MSTIS*, and *MISTIS*. In the examples, the following set of initial imports is assumed:

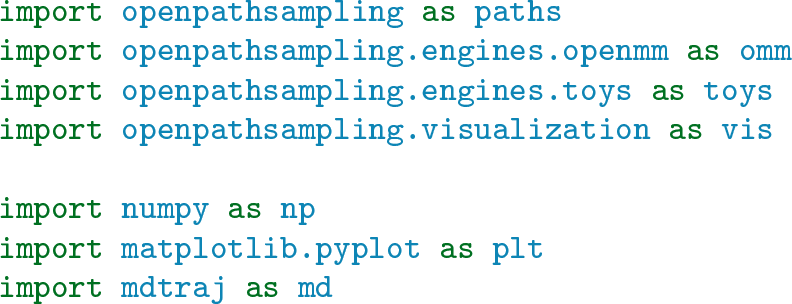

These imports load the required modules, notably the OPS modules, butalso modules such as MDTraj[82], OpenMM [76], the toy dynamics, and the Python plotting modules. We note that the explicit code given inthis section isfor illustrative purposes only, and refers to the 1.0 release. Up-to-date versions of the examples are available as interactive Jupyter notebooks on the website http://openpathsampling.org.

### A. TPS on alanine dipeptide

This example illustrates details about setting up transition path sampling calculations, both with fixed and flexible path length ensembles. This example and the next consider alanine dipeptide (AD) in explicit TIP3P [86] water, using the AMBER96 [87] force field to enable comparison to some previous work [88, 89]. This model has been widely used as a biomolecular test system for rare events methods. We use a VVVR-Langevin integrator at 300*K*[90], with a 2fs timestep and a collision rate of 1 ps^−1^. The long ranged interactions were treated with PME with a cutoff of 1 nm. The AD molecule was solvated with 543 water molecular in a cubic box, and equilibrated at constant pressure of 1 atm using a Monte Carlo barostat. Afterwards the box size was set to the average value of 25.58 Å as obtained in the NPT run. All subsequent simulations were done in the NVT ensemble.

While the example is based on the explicit solvent calculations by Bolhuis, Dellago, and Chandler [91], we differ in several details, including our choice of force field and the details of our ensembles: Ref. 91 used a shorter fixed-length TPS ensemble, whereas we use both a flexible-length TPS ensemble and an 8 ps fixed-length TPS ensemble.

#### 1. Setting up the molecular dynamics

We use OpenMM to set up an MD engine for the AD system. The OpenMM-based OPS engine is essentially a wrapper for the OpenMM `Simulation` object. As with the OpenMM `Simulation`, it requires an OpenMM `System`, and an OpenMM `Integrator`. The interactive OpenMM simulation builder tool [http://builder.openmm.org/] allows us construct an appropriate `System` and `Integrator`. In addition, the OpenMM `Simulation` takes a `properties` dictionary, which we must define.

To build the OPS engine, we also need to fill an options dictionary with some OPS-specific and OpenMM-specific entries. All OPS engines should define `nsteps_per_frame`, the number of time steps per saved trajectory frame, and `n_frames_max`, an absolute maximum trajectory length. For the alanine dipeptide examples, we save every 20 fs (10 steps) and abort the trajectory if it reaches 40 ps:

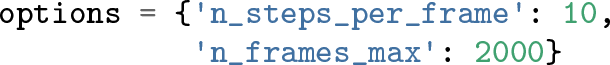

After creating the the OpenMM system, the OpenMM integrator, the OpenMM properties dictionary, and the OPS options dictionary, all of these can be combined to create on OpenMM-based OPS engine:

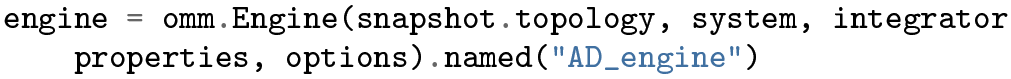

where the `snapshot` is loaded from the PDB with `omm.snapshot_from_pdb(“file.pdb“)`. This command also associates a name with the engine, which make it easier to reload from storage for re-use.

#### 2. Defining states and interfaces

The collective variables of interest for alanine dipeptide are the backbone *ϕ* and *ψ* dihedrals. To create a collective variable for these angles, we use our wrapper around MD-Traj’s `compute_dihedrals` function:

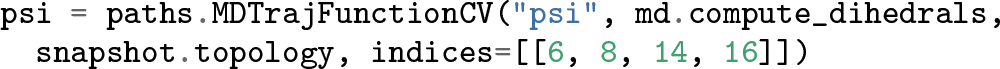

The *ϕ* angle is defined similarly, consisting of the atoms with indices `4, 6, 8`, and `14`.

MDTraj reports dihedral angles in radians. The `MDTrajFunctionCV` wrapper can wrap any function that uses MDTraj; we use the simplest example here for illustrative purposes. It would be straightforward to write a Python function that converts this to degrees, and to use that in place of `md.compute_dihedrals;` the AD MSTIS example in section VI B uses a more complicated approach to wrapping CVs.

In this example, we define two states, *C*_*7eq*_ and *α*_*R*_, similarly to Ref. [91]. Since we are using a different force field, we use slightly different values for the *ψ* angles. Our state *C*_*7eq*_ is defined (in degrees) by 180 ≤ *ϕ* < 0 and 100 ≤ *ψ* < 200 (wrapped periodically), whereas *α*_*R*_ is given by 180 ≤ *ϕ* < 0 and −100 ≤ *ψ* < 0. To convert between degrees and radians, we define deg = np.pi/180. The code to define *C*_*7eq*_ is, and state *α*_*R*_ can be coded accordingly.

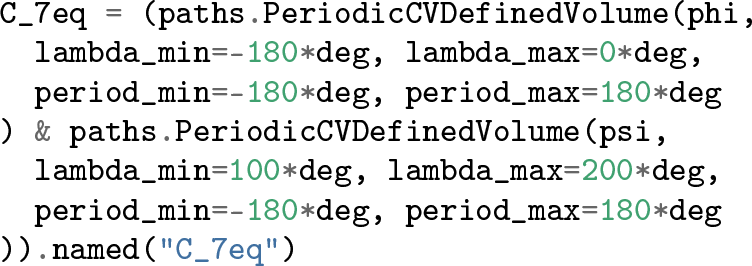

For nonperiodic CVs, the equivalent form is `CVDefinedVolume`, and it does not include the `period_min` and `period_max` arguments. The periodic version allows the *C*_*7eq*_ state to wrap across the periodic boundary in the ψ variable: We define the state from 100 degrees to 200 degrees, even though the function reports values between −180 degrees and 180 degrees. We would get the exact same behavior by setting `lambda_max` to −160 degrees. For a TPS simulation, we only need to define the states — there are no interfaces to define.

#### 3. Setting up the transition network and move scheme

The transition network creates and contains all the ensembles to be sampled. In this case, there is only one ensemble. Later examples will deal with sets of ensembles. The fixed and flexible path length examples diverge here: the fixed path length TPS simulation uses a fixed path length network with path length 400 frames (8 ps), created with

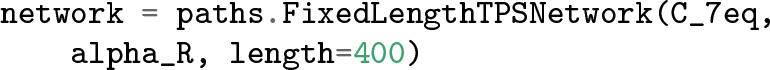

For the flexible path length, which is better in practice, we use:

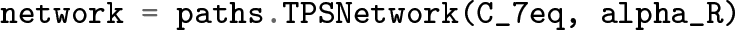

This one line of code selects between the two approaches. Multiple state TPS can beset up similarly. Forinstance, a multiple state TPS with states `A, B`, and `C` (allowing all transitions) can be created by

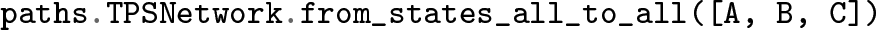

Next, we set up the movescheme. For the TPS simulations, we only need a shooting move. This move scheme is created with

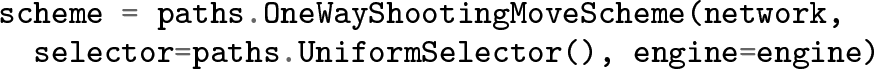

The `selector` defines how to choose the shooting points, e.g., `UniformSelector` selects the points uniformly. Another option would be to use the `GaussianBiasSelector(lambda, alpha, l_0)`, which takes the collective variable lambda and biases the shooting point selection according to 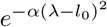, where *l*_0_ is the position of the maximum, and α determines the width ofthe distribution [?].

#### 4. Obtaining initial conditions

We obtained an initial trajectory by running at high temperature (*T* = 500 K) until both states had been visited. Appendix A provides details on this and other possible methods to obtain initial trajectories.

We generate the trajectory for fixed length TPS by taking the appropriate trajectory for flexible length TPS, adding frames to either side, and using the fixed-length ensemble’s `ensemble.split` to select a segment of the appropriate length and satisfying the requirements.

To assign this first trajectory to the ensemble we will be sampling, we use the move scheme’s `scheme.initial_conditions_from_trajectories` method.

#### 5. Equilibration and running the simulation

AllOPS simulation detailsand simulation results are stored in a single NetCDF storage file. The storage requires a `template` snapshot to determine sizes of arrays to save per snapshot. Before running the simulations, we need to create a file to store our results in. A new file named `tps_AD.nc` can be created with

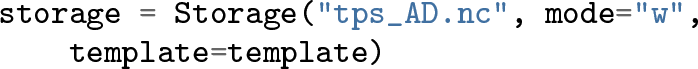

The `PathSampling` simulator object is created with

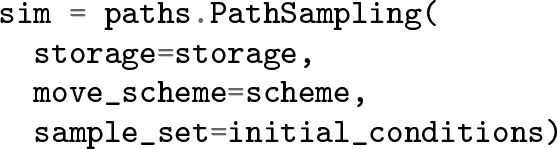

We can run the OPS simulation using with `n_steps` trial moves. We use 10,000 steps for the TPS examples.

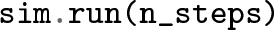

In all molecular simulation approaches initial conditions are unlikely to be representative for the equilibrium distribution (e.g., one could start with the solvent molecules on a grid, orwith a high temperaturesnapshot), and equilibration is usually required before one can take averages of observables. Likewise, we need to equilibrate the path sampling before we can take statistics, when the initial trajectories are not from the real dynamics (e.g., generated with metadynamics or high-temperature simulation). As with MD and MC approaches, the equilibration phase can be just a short version ofthe production run.

#### 6. Analyzing the results

Analysis of a simulation is usually done separately from running the simulation. The first step is to open the storage file with the simulation results will open a file for reading.

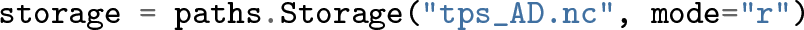

The tables of stored data objects are attributes of the storage. To see the number of items stored, the standard Python len function can be used. For example, `len(storage.steps)` gives the number of Monte Carlo steps run (plus 1 forthe initial conditions).

The move scheme serves as the starting point for much of the analysis. Since there is only one in storage, we obtain the correct move scheme with `scheme = storage.schemes[0]`. The command returns a quick overview ofthe moves performed and information on the acceptance ratios. Since ourTPS movescheme contained only one PathMover, all performed moves were shooting moves. In this example, we find a 56% acceptance ratio for flexible length TPS, and a 50% acceptance rate for fixed length TPS.

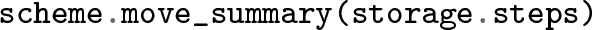

As discussed in Sec. IV, every Monte Carlo `step` in the storage consists of two main parts: the `SampleSet` of active samples, given by `step.active`, and the `PathMoveChange` with details about the move, given by `step.change`. Typically, analysis begins with a loop over steps, and then extracts the relevant information. The first step (step 0) corresponds to the initial conditions. For example, a list of all the path lengths (in frames) can be obtained with which loops over each MC step in `storage.steps`, and takes the length of the trajectory associated with replica ID `0` in the `active` sample set. For TPS, this is the only replica, so this gives us the length of every accepted trajectory, weighted correctly for the ensemble. From here, we can use standard Python libraries to analyze the list, obtaining, for example, the maximum `(max(path_lengths))`, the mean `(np.mean(path_lengths))`, the standard deviation `(np.std(path_lengths))`, or to plot a histogram `(plt.hist(path_lengths))`. In this specific example, we are often interested not in the exact number of frames, but in the time associated with that number of frames. This can be accessed by multiplying the path length by `engine.snapshot_timestep`, which gives the time between saved snapshots. In the case of the OpenMM engine, this result even includes correct units, and we find that the average path length for the flexible path length simulation is 1.6 ps, with a maximum path length of 10.1 ps.

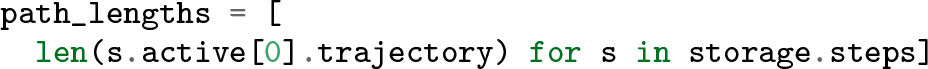

One of the tools for checking the behavior of path sampling simulations, particularly of one-way flexible length path sampling, is the visualization known as the “path tree.” This has several uses, including checking for path decorrelation and that there is sufficient alternation between accepted forward shots and accepted backward shots [92]. In OPS, we generate this object with which works with any list of `steps`, although the visualizations get unwieldy for large numbers of steps. The generator describes how to generate the list of samples to bedisplayed from steps. In TPS, there is only one replica `(replica=0)`, but trees can also be used to track the move history of a specific replica in TIS, where there are multiple replicas.

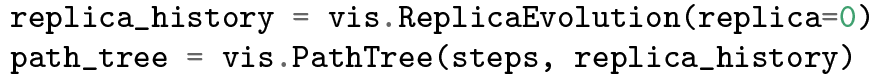

This `PathTree` object only consists of the data and data structures to create and analyze the visualization. The actual image can be generated (in SVG format) using `path_tree.svg()` for visualization in a Jupyter notebook, or written to file. The resulting image, shown in Fig. 7, shows the original trajectory in grey, the forward shots in red, and the backward shots in blue. However, these colors are customizable using CSS options that can be modified by the user. The top uses an additional CSS-based customization to show the individual snapshots. Additional information is shown to the left of the tree. Atthefarleft, a numberindicatesthe MC trial step index. Next to that, a vertical bar contains horizontal lines to indicate groups of correlated paths (paths which share at least on configuration in common).

The list of first-decorrelated paths (the first member of each such group) can be obtained with `replica_history.decorrelated_trajectories`. This number is a good estimate of the number of uncorrelated samples drawn from an ensemble. For the flexible path length simulation, we have 893 decorrelated trajectories, decorrelating on average every 11.2 MC steps. For the fixed path length simulation, there are only 409 decorrelated trajectories, decorrelating every 24.4 MC steps. Note that this set itself has no special relevance, but rather gives an indication of the sampling efficiency.

Besides analyzing the sampling statistics, we can, of course, perform normal MD trajectory analysis on trajectories generated by OPS. For example, suppose we wanted the active trajectory after the 10th MC step. We can obtain this with where, again, TPS only has one replica, with replica ID 0. We can directly analyze this trajectory with the tools in OPS. For example, taking `phi(trajectory)` will give us the list of values of *ϕ* for each frame in the trajectory. Any other OPS collective variable will work similarly, whether it was used in sampling or not.

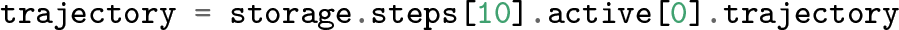

**FIG. 7.**
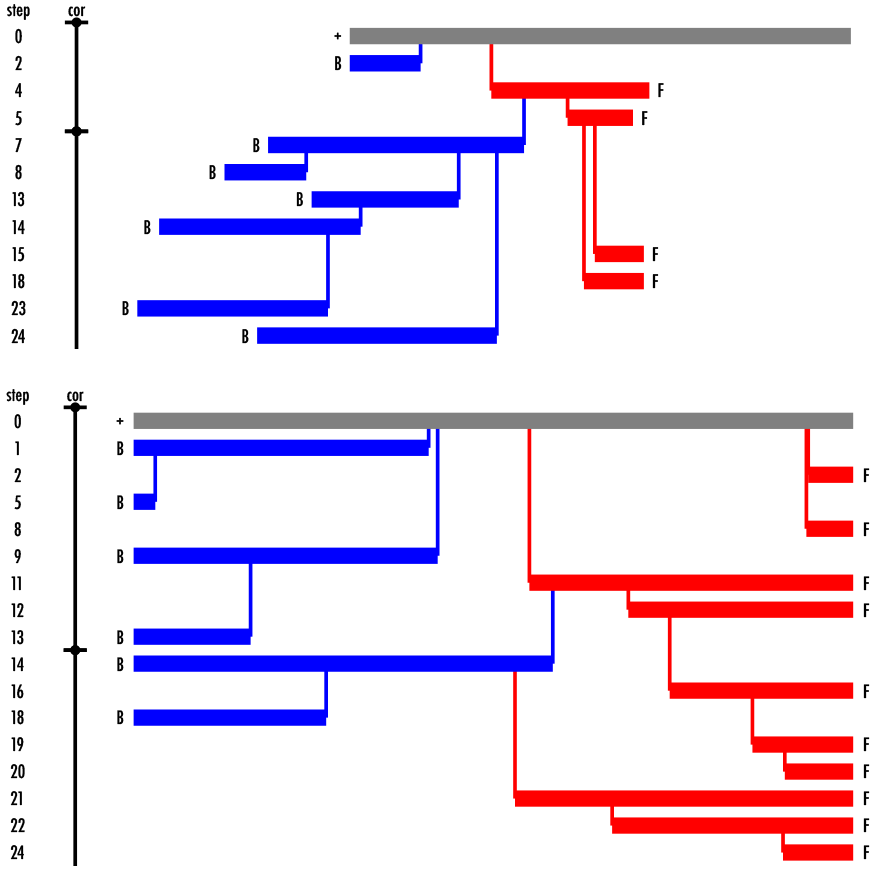
Path sampling history tree for alanine dipeptide TPS simulations from Sec. VI A. Top: The path tree for first 25 trial MC moves using flexible path length TPS. Here the initial path is represented by a grey horizontal line of a length equal to the path length. Going downward, the sequential MC shooting moves are indicated. The dashed vertical line indicates the shooting point. Red and blue horizontal lines indicate forward and backward shots, respectively. Note that these are partial paths, replacing the old path from the shooting point forward (or backwards). The remainder of the path is retained from the previous paths. To the left is indicated the MC step (trial) index. Only accepted paths are shown. The bar to the left indicate complete decorrelation of the previous decorrelated path. Bottom, the path tree for fixed path length TPS. Note that the width is scaled differently; paths in the bottom tree are much longer than the top tree.

The path density gives the numberof paths in the ensemble that visit a particular region in the projected collective variable space. The appropriate histogram requires defining the (inclusive) lower bound of a bin, and the width of the bin in each collective variable. In OPS, we can calculate the path density with utilizing a bin width of 2 degrees.

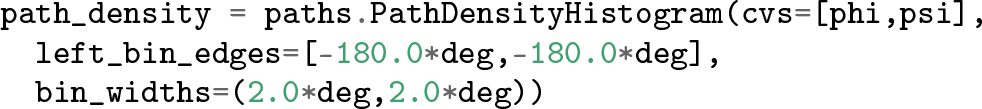

In principle, an OPS path density can be in any numberof collective variables. However, in practice, path densities are almost always shown as 2D projections. Fig. 8 gives the path density for the flexible path length ensemble in the (*ϕ*, *ψ*), plane, along with two representative trajectories.

**FIG. 8.**
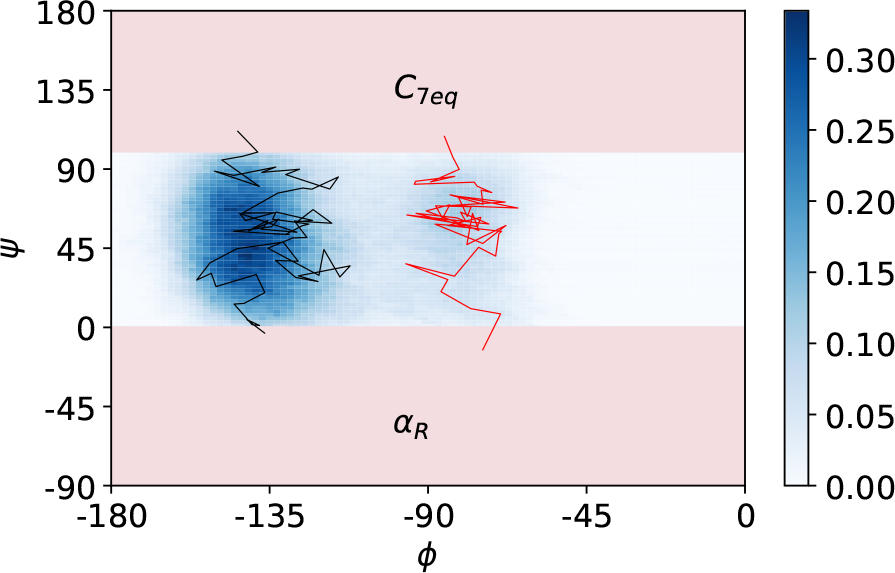
Path density histogram for flexible path TPS ofthe *C*_7eq_ → *α*_*R*_ transition in alanine dipeptide from Sec. VIA. The path density is a 2D histogram ofthe number of paths that traverse a (discrete) ψ – *ϕ* value [66]. On top ofthe path density we plot two individual trajectories, one for each ofthe two observed channels. Note that the left channel between *C*_*7eq*_ and *α*_*R*_ around *φ* ≈ 135 is much more frequently sampled.

If one would rather use other tools, it is possible to convert an OPS trajectory generated by OpenMM to an MDTraj[82] trajectory with

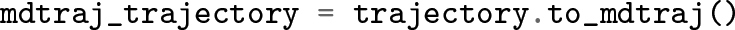

From there, we can analyze the trajectory with MDTraj or export it to any of the file formats supported by MDTraj, to be read in by other analysis programs. In addition, MDTraj can be used as a gateway to other libraries, such as NGLView [93].

The `step.change` starts from the root ofthe move decision tree, and therefore also contains information about what kind of move was decided. This is very simple in TPS, but can be much more complicated forthe move schemes used in TIS. The details that are probably of greatest interest can be accessed with `step.change.canonical`. The nature of a given `step.change.canonical` depends on the type of Monte Carlo move. However, as discussed in Sec. IV and shown in Fig. 2, all `change`s have a few properties: a Boolean as to whether the trial was `accepted`, a link to the actual `mover` that created the change, and a list of attempted samples in `trials`.

Sometimes we might want to study the rejected trajectories, for example, to determine whether they continued to the maximum possible time in flexible length TPS. This could indicate a metastable state that was not considered. The list of rejected samples (which contain the trajectories, as well as the associated ensembles) can be created with

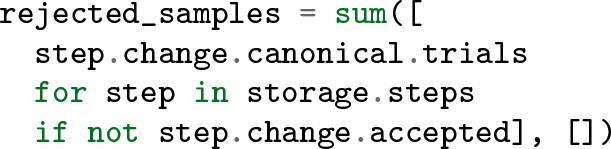

Since each `step.change.canonical.trials` is a list, we use Python’s `sum` function to *add* (extend) the lists with each other. For more complicated move schemes, we might want to add a restriction such as `step.change.mover == desired_mover` with an `and` in the `if` statement. The code above results in a list of Sample objects. The trajectories can be extracted with

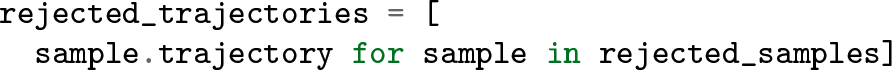

These rejected trajectories can be analyzed in the same way as above.

### B. Multiple state replica exchange TIS on alanine dipeptide

#### 1. Setting up the molecular dynamics

In this example we use the same system as in the previous example, with the same MD engine. In the online Jupyter notebook — which contains additional detail not presented here — we set up the engines from scratch. However, as OPS saves all the details of the engine, we can reload a usable engine from the output file of the previous example. In fact, we can even use that file to reload the collective variables that we defined:

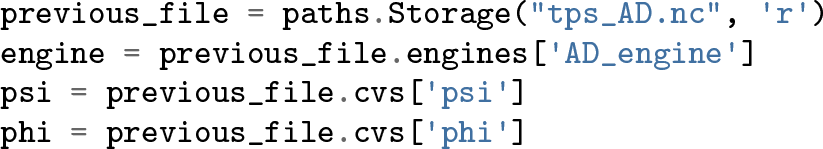

#### 2. Defining states and interfaces

In contrast to the above example we take the MSTIS state definitions for alanine dipeptide from [57] to make the results comparable with that work. The states are defined by a circular region around a center in *ϕ* - *ψ* space, while interfaces are defined by circular regions with increasing diameter *λ*. For instance, for state A we can define:

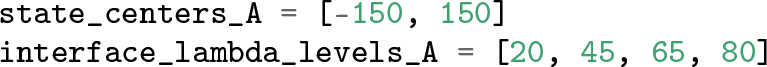

For convenience, Python dict objects can be used to contain the centers and interface levels for all states (e.g., `state_centers[“A“])`, although in this example we will use separate objects for each.

In MSTIS, each state is associated with an order parameter (CV). In simple cases like this one, a single functional can be used for all the order parameters. In OPS, this can be accomplished by creating a single Python function which takes a `Snapshot` as its first argument, and parameters for the functional as the remaining arguments. This was also done implicitly in the previous example, where the `md.compute_dihedrals` Python function is actually a functional with indices as parameters. In this case, we need to explicitly create a Python function with the signature `circle_degree(snapshot, center, cv_1, cv_2)`, where snapshot is an OPS Snapshot, center is a two-member list like `state_centers_A, and cv_1 and cv_2`, are OPS collective variable objects (in all cases, we will use our `phi` and `psi` variables).

In this example, we could redefine the `phi` and `psi` variables inside the function, but using them as parameters has an additional advantage: they will only be calculated once, and then the values will be cached in memory (and optionally saved to disk). This is extremely useful for expensive CVs that are likely to be reused as part of other CVs.

Once the functional has been defined, we can wrap it in an OPS `FunctionCV` for each state. For state *A*:

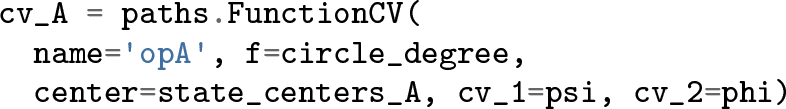

We can now use this CV to define the volume associated with the state:

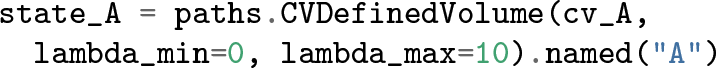

All of this is analogous to the TPS example; however, TIS also requires defining interfaces. These can be created with:

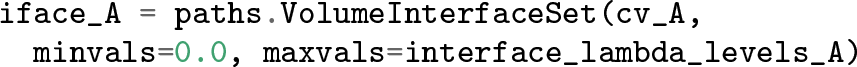

In many cases, the innermost interface volume is identical to the state. For those examples, one could first create the interfaces, and then select the innermost using

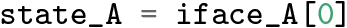

The state definition used here are illustrated in Fig. 5. In this example we restrict the states to {*A*, *B*, *C*, *D*}. The transitions to the E and F states are extremely rare, and requires additional restricted path sampling [57].

#### 3. Setting up the transition network

MSTIS can make use of the optional multistate outer interface, in which all state-to-state paths are allowed, as long as they cross the outer interface `MSOuterInterface`. This special interface allows switching paths from one associated state to another when reversing a transtion path. Note that in all other interface ensembles/replicas such reversal trials are rejected by construction. We create this multi-state outer interface with where the lambdas are the interface levels as defined above, and the dots indicate a short hand for all other state volumes and interfaces.

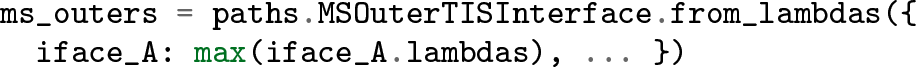

We now construct the `Network` that contains the structure of states and interfaces

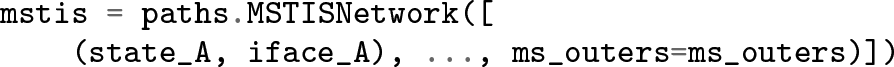

We finally construct the `DefaultScheme` with

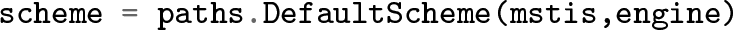

This scheme includes minus moves, moves for the multi-state outer interface ensemble, as well as the standard shooting, path reversal and nearest neighbor replica exchange moves.

**FIG. 9.**
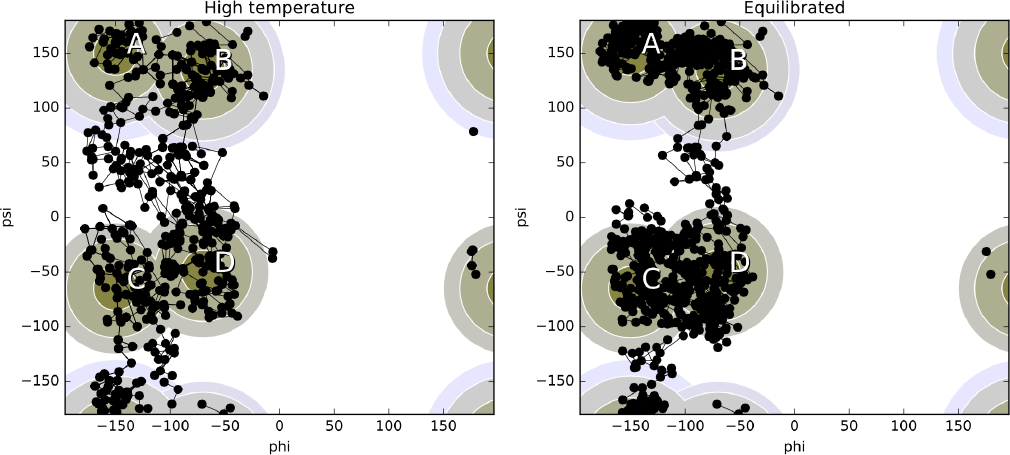
Comparisonbetweeninitial sample after generationat high temperature and room temperature equilibration for alanine dipeptide in the psi-phi plane from Sec. VI B. The trajectories are plotted as connect dots, where each dot represents a snapshot. The stable state and interface definitions for A,B,C,D are plotted in the background. Note that after cooling to room temperature the trajectories are sampling a more narrow path samples, as expected.

#### 4. Obtaining initial conditions

Initial conditionsforthe MSTIS simulation can be obtained with an approach similar to the one used in the TPS example in Sec. VI A. The initial conditions must include a trajectory that satisfies each (interface) ensemble. However, the same trajectory can be reused for multiple ensembles, and an interface that transitions from a given state to another must exit all interfaces associated with the initial state. A trajectory that visits all states has (when considering both the trajectory and its time-reversed version) at least one subtrajectory that represents a transition out of every state.

As the with TPS example, we therefore use the approach described in Appendix A, using a temperature of *T* = 1000 K. Again, the `scheme.initial_conditions_from_trajectories` method is used to identify the specific subtrajectories and associate them with the correct ensembles.

When the initial paths for the minus ensembles are not directly found we use the innermost TIS ensemble trajectories and *extend* them until they match the required (minus) ensemble (or fail in doing so) using the `.extend_sample_from_trajectories` method and associating them with the correct ensembles using `scheme.initial_conditions_from_trajectories`.

#### 5. Equilibrating and running the simulation

As in the TPS examples, the path replicas first need to be equilibrated since the initial trajectories are not from the real dynamics (e.g., generated with metadynamics, high temperature, etc.) and/or because the initial trajectories are not likely representatives ofthe path ensemble (e.g., if state-to-state transition trajectories are used for all interfaces).

As with straightforward MD simulations, running equilibration can be the same process as running the total simulation. However, in path sampling we could equilibrate without replica exchange moves or path reversal moves, for instance. In the example below, we create a new ‘MoveScheme’ that only includes shooting movers, to achieve equilibration of the interface ensemble replicas, and run this scheme for 500 steps the way we run any other scheme, using the `PathSampling` object.

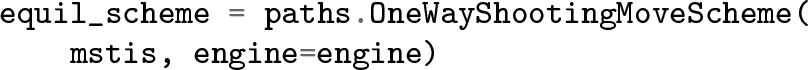

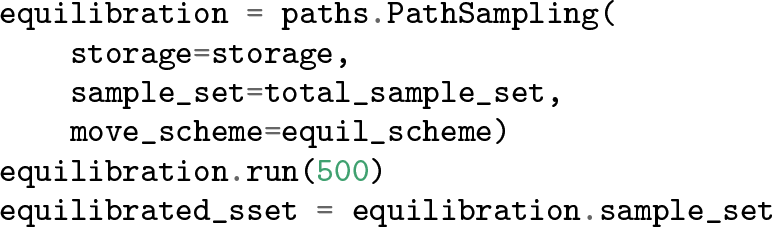

**FIG. 10.**
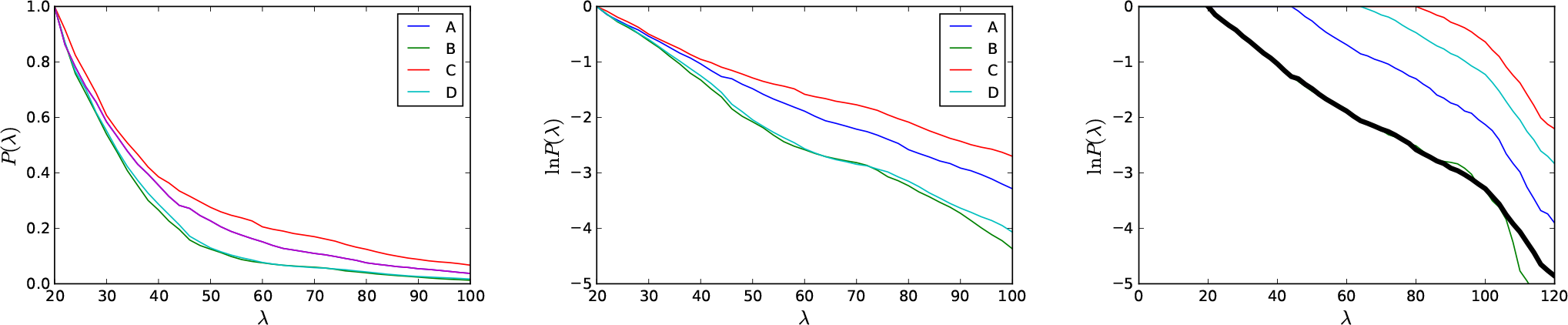
TIS Crossing probabilities for alanine dipeptide, from section VI B. *Left:* Total crossing probability as function of the order parameter (CV)*λ* for each individual state (A-D). *Center:* Natural logarithm of the total crossing probability per state. *Right:* Per interface crossing probabilities for state A. The master curve (black) is obtained by reweighting [60, 69].

Figure 9 shows the set of initial and final samples. Note that the large coverage of phase space at high temperature narrows after cooling down, as expected.

Fianlly, we run the simulation for 100,000 steps using the PathSampling object as in previous example, with the default scheme as defined above, a `Storage` object, and the initial conditions from the equilibration.

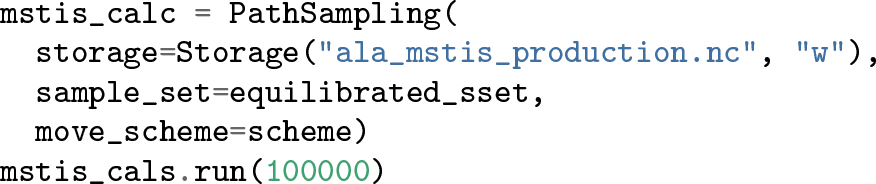

#### 6. Analyzing the results

To do analysis on the path simulation results we first have to load the production file for analysis

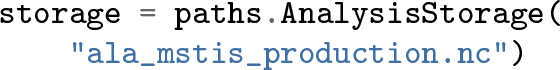

Then we can run analysis on this storage. One of the main objectives for doing multiple state replica exchange TIS is to compute the rate constant matrix. To obtain the rate constant matrix, we run which gives as an output the full rate constant matrix *k*_*IJ*_, obtained from a computation of the fluxes *ϕ*_*0I*_, the crossing probabilities *P*_*I*_(*λ*_*mI*_|*λ*_0*I*_) and the conditional transition matrix *P*_*I*_(*λ*_0*J*_|*λ*_*mI*_) (see Eq.5).

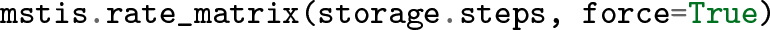

**TABLE II.** Rate constant matrix for alanine dipeptide. The average rate constant matrix forthe four-state Markov model based on several independent runs. Rows denote leaving, columns arriving states. Subscript denotes error in the last 2 digits.

|  | A | B | C | D |
| --- | --- | --- | --- | --- |
| A | 0 | 0.065<sub>12</sub> | 0.0035<sub>14</sub> | 0.00097<sub>25</sub> |
| B | 0.076<sub>06</sub> | 0 | 0.0012<sub>02</sub> | 0.0011<sub>07</sub> |
| C | 0.053<sub>11</sub> | 0.011<sub>03</sub> | 0 | 0.12<sub>02</sub> |
| D | 0.011<sub>02</sub> | 0.0056<sub>24</sub> | 0.069<sub>09</sub> | 0 |

**TABLE III.**
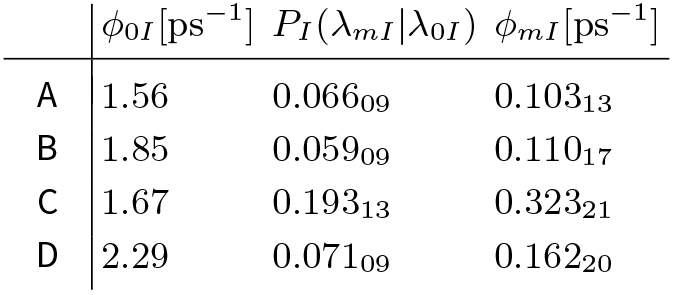
Fluxes and outer interface crossing probabilities for TIS simulation of alanine dipeptide. Flux at the first interface (second column), the crossing probability from the first to the outermost interface (third column) and the flux at the outermost interface (last column).

**TABLE IV.** Conditional transition probability matrix between alanine dipeptide states. These probabilities follow directly from the path sampling in the multistate outer ensemble. Rows denote leaving, columns arriving states.

|  | A | B | C | D |
| --- | --- | --- | --- | --- |
| A | 0.35<sub>06</sub> | 0.61<sub>06</sub> | 0.032<sub>08</sub> | 0.0090<sub>12</sub> |
| B | 0.72<sub>03</sub> | 0.26<sub>03</sub> | 0.011<sub>02</sub> | 0.011<sub>08</sub> |
| C | 0.17<sub>02</sub> | 0.033<sub>07</sub> | 0.40<sub>03</sub> | 0.40<sub>01</sub> |
| D | 0.066<sub>05</sub> | 0.034<sub>13</sub> | 0.42<sub>03</sub> | 0.48<sub>04</sub> |

An example of such a rate constant matrix computation is shown in Table II which agrees well with the results in Ref. [57]. For comparison, the computed fluxes *ϕ*_0*I*_ and crossing probabilities *P*_*I*_(*λ*_*mI*_|*λ*_0*I*_) are presented in Table III, while Table IV reports the conditional state-to-state transition matrix *P*_*I*_(*λ*_*0J*_|*λ*_*mI*_), which represents the probabilities to reach a state provided that the trajectories have passed the outermost interface of a state.

OPS also includes other analysis tools such as the crossing probabilities and the sampling statistics. Both are important for purposes of check the validity of the simulations results. The crossing probability graphs in Fig. 10 can be helpful in interpreting the rate matrix. The sampling statistics provides the Monte Carlo acceptance ratio for the different movers. Of course, each trajectory in the ensemble can accessed and scrutinized individually, as in previous sections.

### C. Example: MISTIS on a three-state 2D model system

This example deals with a three-state 2D model system, which we also refer to as a *toy model*. OPS includes simple code to simulate the dynamics of small toy models like the one considered here. This is intended for use for either educational purposes or for rapid prototyping of new methodologies. Since the overall path sampling code is independent of the underlying engine, many types of new methods could be developed and tested on the toy models and would be immediately usable for more complicated systems, simply by changing the engine.

#### 1. Setting up the molecular dynamics

We create a simple 2D model with a potential consisting of a sum of Gaussian wells:

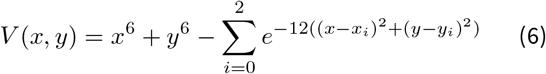

with (*x*_0_, *y*_0_) = (−0:5, 0:5), (*x*_1_, *y*_1_) = (−0:5, −0:5), and (*x*_2_, *y*_2_) = (0:5, −0:5) using

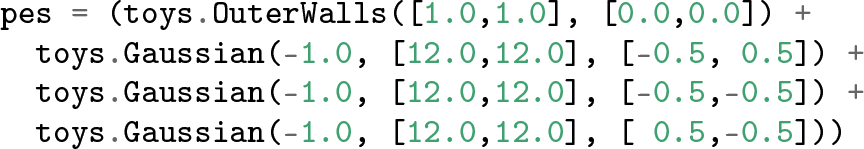

This results in a potential energy surface with three stable states, caused by the Gaussian wells at (−0.5, −0.5), (0.5, −0.5), and (−0.5,0.5). We call those states `A, B,` and `C`, respectively. This potential interface surface, along with the state and interface definitions described below, is illustrated in Fig. 11.

To integrate the equations of motion, we use the BAOAB Langevin integrator of Leimkuhler and Matthews [94], which we initialize with

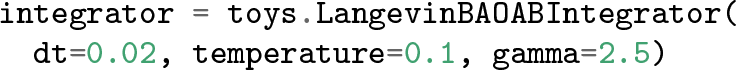

**FIG. 11.**
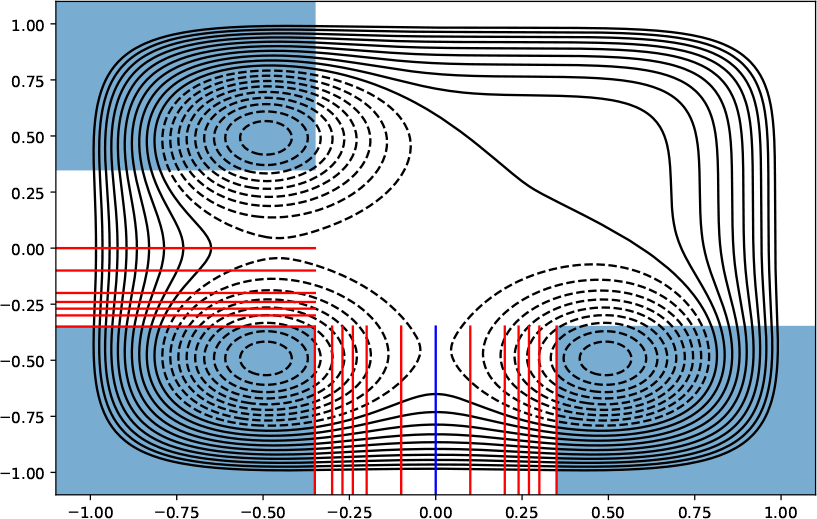
Potential energy surface, states, and interfaces for the 2D toy model. States are light blue, boundaries of normal interfaces are red, and the boundary ofthe multiple state outer interface is dark blue. For clarity when showing multiple interface sets, the interface bondaries are only drawn part ofthe way. They continue in an infinite straight line.

The toy engine employs units where *k*_*B*_ = 1. The “topology” forthe toy engine stores the number of spatial degrees of freedom, as well as a mass for each degree of freedom and the potential energy surface. We also create an options dictionary forthe engine.

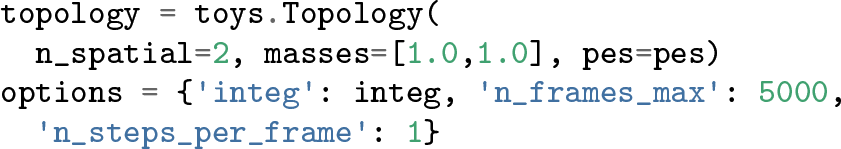

We then instantiate an engine with `toy_engine = toys.Engine(options, topology)`.

#### 2. Defining states and interfaces

In this calculation, we will set up multiple interface set transition interface sampling (MISTIS)[71]. This involves defining different interface sets for each transition. To simplify, and to highlight some ofthe flexibility of MISTIS, we will only focus on the *A* → *B*, *B* → *A*, and *A* → *C* transitions.

First, we define simple collective variables. We use the Cartesian *x* and *y* directions for both our state definitions and for our interfaces. We define these as standard Python functions, for example and `xval` is similar, but returns the `[0][0]` element, where the first index refers to the atom number and the second to the spatial dimension. We wrap functions these in OPS collective variables with, for example, `cvX = FunctionCV(“x”, xval)`. We’ll define a volume called `x_lower` for *x* < −0.35 with `CVDefinedVolume(cvX, float(“-inf”), −0.35)`. Similarly we define `x_upper` for *x* ≥ 0.35, and use the same bounds for *y* with `y_lower` and `y_upper`. With these, we define our states as

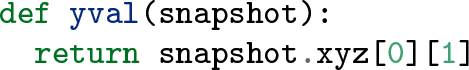

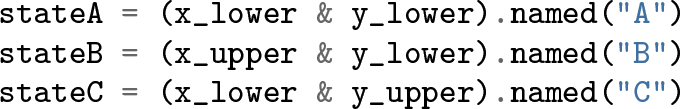

For our TIS analysis, the order parameter must increase with the interface. So forthe *B* → *A* transition we create anothei collective variable, `cvNegX`, based on a function that return; `-snapshot[0][0]`. For all of these, we will set interfaces al −0.35, −0.3, −0.27, −0.24, −0.2, and −0.1. The *A* → *C* transition has an additional interface at 0.0 The *A* → *B* and *B* → *A* transitions will share a multiple state outer interface at 0.0. Note that, for the *B* → *A* transition, the interface associated with `cvNegX` = −0.35 is actually at *x* = 0.35 since `cvNegX` returns −*x*. These interfaces are created by, fo example: with similar lines for `interfacesAC` and `interfacesBA`.The multiple state outer interface, which connects the *A* → *B* and *B* → *A* transitions, can be created at *x* = 0.0 with

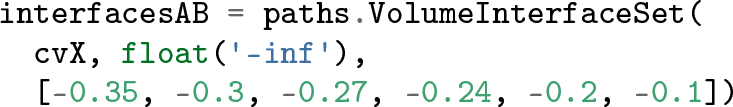

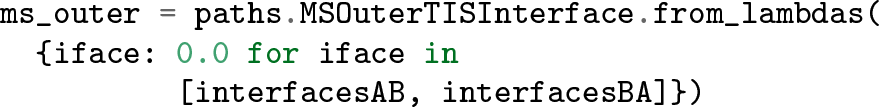

#### 3. Setting up the transition network and move scheme

Like the `MSTISNetwork`, the syntax for setting up a `MISTISNetwork` requires a list of tuples. However, since MISTIS requires a *final* state as well an initial state, it also requires that the final state be included as an extra piece of information in that tuple. So we set up our desired MISTIS network with

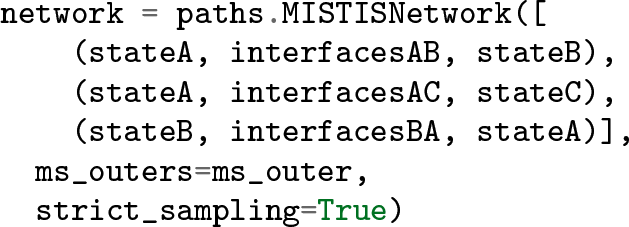

The `strict_sampling` argument means that an *A* → *C* path will be rejected if sampling the *A* → *B* transition. Note that *A* is the initial state for transitions to two states, whereas *B* is the initial state for transitions to one state, and *C* is not an initial state at all. The flexibility to define arbitrary reaction networks is an important aspect of the MISTIS approach.

The move scheme is set up in exactly the same way as for MSTIS: `scheme = DefaultScheme(network, engine)`. One could also use a single replica move scheme with a MISTIS network, just as was done in the MSTIS example.

#### 4. Obtaining initial conditions

Here, we will use the bootstrapping approach to obtain initial trajectories. This approach is most effective with simple systems like this, where the collective variables we have chosen as order parameters are good representations of the actual reaction coordinate. The bootstrapping runs separately on each transition *A* → *B*, *A* → *C*, and *B* → *A*. Given an initial snapshot `snapA` in state *A*, the initial samples forthe *A* → *B* transition can be obtained with

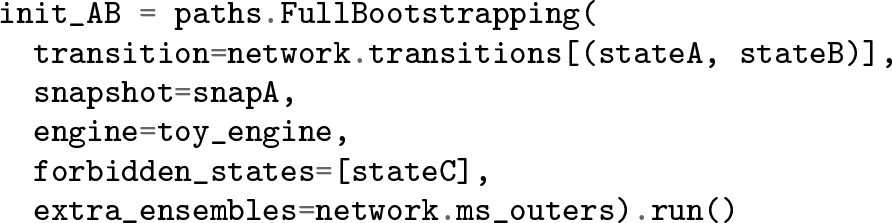

This will create a trajectory for each of the normal interface ensembles, as well as the multiple state outer interface ensembles. The other transitions can be prepared similarly, although they can omit the `extra_ensembles` option, since there is only one multiple state outer ensemble to fill. Whereas snapshots for the OpenMM engine used in the previous examples came from PDBs or other files, for the toy engine, the initial snapshot can be manually created:

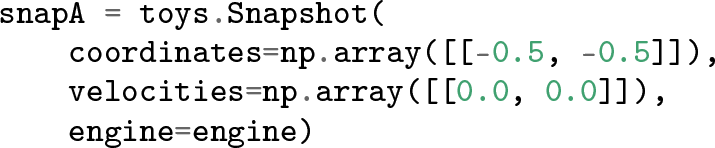

The individual sample sets created by the `FullBootstrapping` approach can be combined into one using

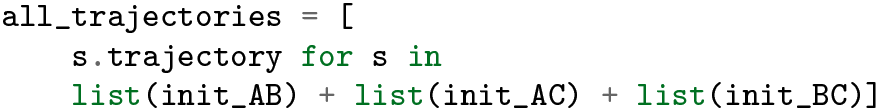

From here, the setup follows that of the MSTIS example: the trajectories can be assigned to ensembles using `scheme.initial_conditions_from_trajectories`, and the minus ensembles, of which there is one for each state, can be filled using `minus.extend_sample_from_trajectories`.

#### 5. Running the simulation and analyzing the results

The path sampling follows exactly as with the previous examples. The `PathSampling` object is created with a storage file, the move scheme, and the initial conditions. We use the `.run(n_steps)` method to run the simulation.

One difference with the MSTIS approach is that trajectories from the multiple set minus interface cannot be used to calculate the flux. To obtain the flux, we do a separate calculation, which we call `DirectSimulation`, and which runs a molecular dynamics trajectory and calculates the flux and the rates from the direct MD.

Setting up the `DirectSimulation` requires the same `toy_engine` object. The set of all states if given by

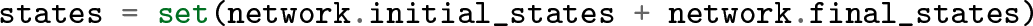

To determine the flux out of a given state and through a given interface, it needs the pairs of (state, interface) for each transition. We can create this with

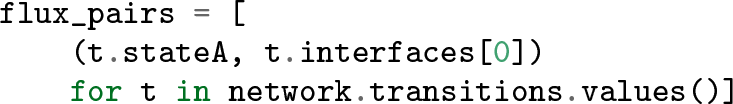

**TABLE V.** **Rates constants for the the toy model**, multiplied by 10^4^. Subscripts indicate error in the last two digits. Left: Rate constant calculated from a very long direct molecular dynamics simulation. Right: Rate constant calculated using MISTIS. By symmetry, all three rate constants in each table should be nearly the same.

|  | A | B | C |  | A | B | C |
| --- | --- | --- | --- | --- | --- | --- | --- |
| A | — | 1.98<sub>14</sub> | 1.95<sub>17</sub> | A | — | 1.94<sub>43</sub> | 2.20<sub>65</sub> |
| B | 2.00<sub>16</sub> | — | — | B | 2.10<sub>37</sub> | — | — |

The simulation is then created with where we choose not to store the output, and where we can use any snapshot as our initial snapshot. The method `sim.run(n_steps)` runs the simulation for the given number of MD steps. Note that, although the direct simulation here is forthe MISTIS network, it would work equally well for any other network. However, the flux calculation based on the minus interface is more convenientforthe MSTIS case.

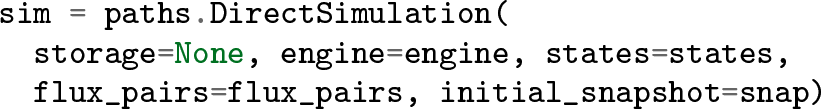

Once the direct simulation has been run, we can obtain the flux from it using `sim.fluxes`. This returns a Python dictionary with the (`state, interface`) pairs as keys and the calculated flux as value. Prior to the rate matrix calculation, we can set the fluxes for the network by using `network.set_fluxes(sim.fluxes)`.

Aside from setting the flux, the analysis forthe MISTIS network is exactly as it is for other path sampling methods. The rate matrix for this model is presented in Table V. Note that the rate matrix only includes the specific transitions we selected forstudy by MISTIS; others are not listed. The MISTIS rates represent the average of 10 runs of 10^5^ MC steps each, with the standard deviation as the reported error. To demonstrate correctness, we compare these rates to those from a direct MD simulation (also performed using OPS), with a length of 8 × 10^8^ frames. The cumulative MD time for the 10 MISTIS simulations was less than 1.8 × 10^7^ frames. Errors forthe direct MD rate were determined by splitting the total simulation into 10 sequential blocks and calculating the standard deviation of the rate in each block. Rates and error bars from the two methods compare favorably, even though the MD simulation took more than 40 times more CPU time.

## VII. CONCLUSION

In this paper we have presented a new easy-to-use Python framework for performing transition path sampling simulations of (bio)molecular systems. The OpenPathSampling framework is extensible and allows for the exploration of new path sampling algorithms by building on a variety of basic operations. As the framework provides a simple abstraction layer to isolate path sampling from the underlying molecularsimulation engine, new molecularsimulation packages can easily be added. Besides being able to execute existing complex path sampling simulations schemes, tools are provided to facilitate the implementation of new path sampling schemes built on basic path sampling components. In addtion, tools for analaysis of e.g., rate constants, are also provided. Modules that provide additional functionality are continuously added to be used by the community (see, e.g., the repositories at https://gitlab.e-cam2020.eu/Classical-MD_openpathsampling).

In summary, the OpenPathSampling package can assist in making the tranistion path sampling approach easierto use forthe (bio)molecularsimulation community.

## ACKNOWLEDGMENTS

DWHS and PGB acknowledge support from the European Union’s Horizon 2020 research and innovation program, under grant agreement No 676531 (project E-CAM). JDC acknowledges support from Cycle for Survival, NIH grant P30 CA008748, and NIH grant R01 GM121505. JDC, JHP, and DWHS gratefully acknowledge support from the Sloan Kettering Institute. FN acknowledges ERC consolidator grant 772230 “ScaleCell”, DFG NO 825/3-1, and SFB1114, project A04.

The authors are grateful for feedback from many people who helped beta-test the software, whose names are listed at http://openpathsampling.org/latest/acknowledgments.html. The authors are particularly grateful to Sander Roet (University of Amsterdam) for his feedback, and to Jocelyne Vreede (University of Amsterdam) for the feedback obtained by using OPS asa teaching tool in courses on biomolecularsimulation.

## Appendix A: Obtaining an initial trajectory

Just as configurational Monte Carlo requires a valid initial configuration for input, path sampling Monte Carlo requires a valid initial trajectory for input. And just as with configurational Monte Carlo, an unrealistic initial state can equilibrate into a realistic state, but more realistic starting conditions are preferred. Unrealistic starting conditions can take longer to equilibrate, and can get trapped in unrealistic metastable basins. In the case of path sampling, this can mean sampling a transition with a much higher energy barrier than is realistic.

Obtaining a good first trajectory is thus of paramount importance. However, there is no single best method to do so. Here we review a few options, and explain how OPS can facilitate first trajectory generation. In all of these, the key OPS functions that simplify theprocess are the `Ensemble.split` function, which can identify subtrajectories that satisfy the desired ensemble, and the `MoveScheme.initial_conditions_from_trajectories` function, which attempts to create initial conditions for the desired move scheme bsaed on given trajectories. The fundamental trade-off forthese approaches is between how “realistic” the initial trajectory is, and how computationally expensive it is to obtain the first trajectory.

## 1. Long-time MD

While a transition from an unbiased MD trajectory will provide a realistic initial trajectory, these are difficult to obtain for rare events. Nevertheless, distributed computing projects like Folding@Home [95] and special-purpose computers for MD such as Anton [96], might yield trajectories that include a transition. The `Ensemble.split` function can then select subtrajectories that satisfy a desired ensemble.

In these trajectories frames are often saved very infrequently, leading to transitions with onlyoneortwo (oreven zero) frames in the “no-man’s land” between the states. Path sampling requires at least a few to a few tens of frames. A `CommittorSimulation` using the desired states as endpoints and any frames between the two states as input could generate an initial trajectoriy by joining two path segments ending in different states, provided the committor for at least one ofthe intermediate frames is reasonable.

## 2. High temperature MD

In the alanine dipeptide example, we use MD at high temperature to increase the probability of getting a transition. This method could cause problems in larger systems, by allowing transitions that are not accessible at the relevant low temperature. However, it works well on simple systems such asAD,and is very easy to set up.

First, we create an engine with a higher temperature. For the AD example, we use OPS’s OpenMM engine. For the 2-state system, we used a high temperature of 500*K*. Forthe 4-state system, we used a high temperature of 1000*K* in order to easily reach the higher-lying states.

To ensure that we visit all states, we generate a trajectory usingthe ensemble which createsthe union of `AllOutXEnsembles` for each state. Running with this ensemble as the continue condition means that the trajectory will stop with the first trajectory that does not satisfy it, i.e. the first trajectory with at least one frame in each state. For MSTIS, this guarantees that a subtrajectory (or its reversed version) will exist for every path ensemble.

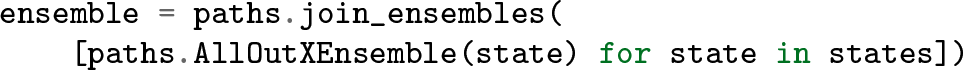

We obtain the relevant trajectories by using the `ensemble.split` method with the outermost ensemble for each sampling transition (See also Ref. [45]).

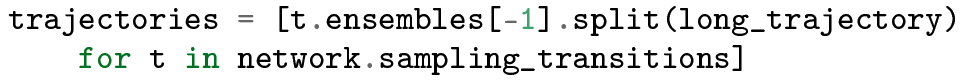

These trajectories can then be given to the `MoveScheme.initial_conditions_from_trajectories` method.

Relaxing the high temperature initial trajectories down to ambient conditions might be difficult, requiring many shooting attempts before a valid room temperature path is created. Again, a `CommittorSimulation` can alleviate this problem, by joining a forward and backward committorsegmentsfrom a snapshot with a finite committor value.

## 3. Bootstrapping/Ratcheting

In the toy model example in Sec. VI C 4, we use a “bootstrapping” approach, which is specifically useful forTIS. In this approach, we initialize a trajectory in a stable state, e.g. A, and perform MD until the first interface is crossed, which allows the first interface ensemble to be populated,. Subsequently, TIS is performed until the second interface is crossed, allowing the second interface to be populated, etc etc. In this way one can ratchet oneself up the barrier and populate each TIS interface. All this is taken care of by the `FullBootstrapping` method. Note that this path ensemble needs to be equilibrated subsequently.

## 4. Using biased trajectories

The use of unbiased dynamics is not necessary, as the goal is to obtain an initial trajectory that is just “reasonably close” to the unbiased dynamics. Subsequent path sampling will then equilibrate the trajectories with the unbiased dynamics.

Recent work [51] has employed metadynamics [15] to obtain an initial trajectory. Although metadynamics biases the underlying dynamics, the first transition in a metadynamics simulation will not have added much bias to the barrier region. Therefore, further path sampling can equilibrate a first metadynamics trajectories into the unbiased dynamics path ensemble. This initial metadynamics trajectory could be generated with PLUMED [80], and then read into OPS.

The same basic approach could be employed for other approaches to generate a non-physical initial transition trajectory, including steered MD [97], nudged elastic band [98], or the string method[99].

## References

[1] I. Buch, T. Giorgino, and G. De Fabritiis, Proc. Nat. Acad. Sci. USA 108, 10184 (2011).

[2] N. Plattner, S. Doerr, G. D. Fabritiis, and F. Noe, Nature Chem-. istry 9, 1005 (2017).

[3] D.-A. Silva, G. R. Bowman, A. Sosa-Peinado, and X. Huang, PLoS Computational Biology 7, e1002054 (2011).

[4] C. Schütte, A. Fischer, W. Huisinga, and P. Deuflhard, Journal of Computational Physics 151,146 (1999).

[5] C. Schütte and W. Huisinga, in Handbook of Numerical Analysis, edited by P. G. Ciaret, and J.-L. Lions (Elsevier, ADDRESS, 2003), Vol. X, pp. 699–744.

[6] F. Noé, I. Horenko, C. Schütte, and J. C. Smith, J. Chem. Phys. 126, 155102 (2006).

[7] J. D. Chodera, N. Singhal,V. S. Pande, K. A. Dill, and W. C. Swope, J. Chem. Phys. 126, 155101 (2007).

[8] D. Chandler, in Classical and Quantum Dynamics in Condensed Phase Simulations, edited by B. J. Berne, G. Ciccotti, and D. F. Coker, (World Scientific, ADDRESS, 1998), Chap. Barrier crossings: classical theory of rare but important events, pp. 3–23.

[9] P. G. Bolhuis, D. Chandler, C. Dellago, and P. Geissler, Ann. Rev. Phys. Chem. 53,291 (2002).

[10] G. M. Torrie, and J. P. Valleau, Chem. Phys. Lett. 28, 578 (1974).

[11] E. Carter, G. Ciccotti, J. T. Hynes, and R. Kapral, Chem. Phys. Lett. 156,472 (1989).

[12] T. Huber, A. Torda, W. van Gunsteren, J. Comput. Aided Mol. Des. 8, 695 (1994).

[13] H. Grubmüller, Phys. Rev. E 52, 2893 (1995).

[14] A. F. Voter, J. Chem. Phys. 106,4665 (1997).

[15] A. Laio and M. Parrinello, Proc. Nat. Acad. Sci. USA 99,12562 (2002).

[16] E. Darve and A. Pohorille, J. Chem. Phys. 115, 9169 (2001).

[17] Y. Sugita, and Y. Okamoto, Chem. Phys. Lett. 314, 141 (1999).

[18] E. Marinari and G. Parisi, Europhys. Lett. 19, 451 (1992).

[19] L. Zheng, M. Chen, and W. Yang, Proceedings of the National Academy of Sciences 105, 20227 (2008).

[20] Y. Q. Gao, J. Chem. Phys. 128, 064105 (2008).

[21] C. Dellago, P. G. Bolhuis, F. S. Csajka, and D. Chandler, J. Chem. Phys. 108,1964 (1998).

[22] C. Dellago, P. G. Bolhuis, and P. L. Geissler, Adv. Chem. Phys. 123, 1 (2002).

[23] C. Dellago and P. G. Bolhuis, Adv Polym Sci 221,167 (2009).

[24] R. Allen, D. Frenkel, and P. ten Wolde, J. Chem. Phys. 124, xxx (2006).

[25] F. Cerou, A. Guyader, T. Lelievre, and D. Pommier, J. Chem. Phys. 134, xx (2011).

[26] A. K. Faradjian, and R. Elber, J. Chem. Phys. 120,10880 (2004).

[27] M. Villen-Altamirano and J. Villen-Altamirano, Eur. Trans. Telecom. 13,373 (2002).

[28] J. T. Berryman, and T. Schilling, J. Chem. Phys. 133, 244101 (2010).

[29] A. Dickson, A. Warmflash, and A. R. Dinner, J. Chem. Phys. 131, 154104 (2009).

[30] G. Huber and S. Kim, Biophysical Journal 70, 97 (1996).

[31] B. W. Zhang, D. Jasnow, and D. M. Zuckerman, The Journal of Chemical Physics 132, 054107 (2010).

[32] J.-H. Prinz, H. Wu, M. Sarich, B. Keller, M. Senne, M. Held, J. D. Chodera, C. Schütte, and F. Noé, J. Chem. Phys. 134,174105 (2011).

[33] N. Singhal, C. D. Snow, and V. S. Pande, J. Chem. Phys. 121,415 (2004).

[34] J. Rogal and P. G. Bolhuis, J. Chem. Phys. 129, 224107 (2008).

[35] S. Pronk, G. R. Bowman, B. Hess, P. Larsson, I. S. Haque, V. S. Pande, I. Pouya, K. Beauchamp, P. M. Kasson, and E. Lindahl, in 2011 International Conference for High Performance Computing, Networking, Storage and Analysis (SC) (PUBLISHER, ADDRESS, 2011), pp. 1–10.

[36] J. Preto and C. Clementi, Phys. Chem. Chem. Phys. 16,19181 (2014).

[37] V. Balasubramanian, I. Bethune, A. Shkurti, E. Breitmoser, E. Hruska, C. Clementi, C. Laughton, and S. Jha, in 2016 IEEE 12th International Conference on e-Science (e-Science) (PUBLISHER, ADDRESS, 2016), pp. 361–370.

[38] H. Wu, A. S. J. S. Mey, E. Rosta, and F. Noé, The Journal of Chemical Physics 141, 214106 (2014).

[39] H. Wu, F. Paul, C. Wehmeyer, and F. Noé, Proceedings of the National Academy of Sciences 113, E3221 (2016).

[40] A. S. J. S. Mey, H. Wu, and F. Noe, Phys. Rev. X 4, 041018 (2014).

[41] E. Rosta and G. Hummer, J. Chem. Theory Comput 11, 276 (2015).

[42] P. Eastman and V. S. Pande, Computing in Science and Engineering 12,34 (2010).

[43] P. Eastman, M. Friedrichs, J. D. Chodera, R. Radmer, C. Bruns, J. Ku, K. Beauchamp, T. J. Lane, L.-P. Wang, D. Shukla, T. Tye, M. Houston, T. Stitch, and C. Klein, J. Chem. Theor. Comput. 9, 461 (2012).

[44] A. Lervik, E. Riccardi, and T. S. van Erp, Journal of Computational Chemistry 38,2439 (2017).

[45] D. W. H. Swenson, J.-H. Prinz, J. Chodera, and P. G. Bolhuis, to be published (2018).

[46] P. G. Bolhuis, and C. Dellago, Reviews of Computational Chemistry (Wiley-VCH, Hoboken, 2009).

[47] E. Guarnera and E. Vanden-Eijnden, J. Chem. Phys. 145,024102 (2016).

[48] J. Juraszek and P. G. Bolhuis, Proc. Nat. Acad. Sci. USA 103, 15859 (2006).

[49] M. Grünwald, C. Dellago, and P. L. Geissler, J. Chem. Phys. 129, 194101 (2008).

[50] R. G. Mullen, J.-E. Shea, and B. Peters, J. Comput. Theory Chem. 11, 2421 (2008).

[51] Z. F. Brotzakis, and P. G. Bolhuis, J. Chem. Phys. 145, 164112 (2016).

[52] T. S. van Erp, D. Moroni, and P. G. Bolhuis, J. Chem. Phys. 118, 7762 (2003).

[53] J. Rogal, W. Lechner, J. Juraszek, B. Ensing, and P. G. Bolhuis, J. Chem. Phys. 133,174109 (2010).

[54] C. Dellago, P. G. Bolhuis, and D. Chandler, The Journal of Chemical Physics 110,6617(1999).

[55] G. M. Torrie, and J. P. Valleau, Chem. Phys. Lett. 28, 578 (1974).

[56] T. S. van Erp, and P. G. Bolhuis, J. Comput. Phys. 205,157 (2005).

[57] W.-N. Du, K. A. Marino, and P. G. Bolhuis, J. Chem. Phys. 135, 145102 (2011).

[58] W. Du and P. G. Bolhuis, J. Chem. Phys. 140,195102 (2014).

[59] F. Noe, D. Krachtus, J. Smith, and S. Fischer, J. Chem. Theor. Comput. 2, 840 (2006).

[60] D. Minh and J. Chodera, JCP 131, 134110 (2009).

[61] W.-N. Du and P. G. Bolhuis, J. Chem. Phys. 139, 044105 (2013).

[62] N. G. van Kampen, Stochastic processes in physics and chemistry, 2nd ed. (Elsevier, ADDRESS, 1997).

[63] F. Noe, C. Schutte, E. Vanden-Eijnden, L. Reich, and T. R. Weikl, Proceedings of the National Academy of Sciences 106, 19011 (2009).

[64] P. G. Bolhuis, and C. Dellago Eur. Phys. J. ST 224, 2409 (2015).

[65] P. Bolhuis, Proc. Nat. Acad. Sci. USA 100,12129 (2003).

[66] J. Vreede, J. Juraszek, and P. G. Bolhuis, Proc. Nat. Acad. Sci. USA 107, 2397 (2010).

[67] Z. F. Brotzakis, and P. G. Bolhuis, to be published (2017).

[68] T. van Erp, Phys. Rev. Lett. 98, 268301 (2007).

[69] P. G. Bolhuis, J. Chem. Phys. 129, 114108 (2008).

[70] T. S. van Erp, Adv. Chem. Phys. 151, 27 (2012).

[71] D. W. H. Swenson and P. G. Bolhuis, J. Chem. Phys. 141, 044101 (2014).

[72] R. Cabriolu, K. M. S. Refsnes, P. G. Bolhuis, and T. S. van Erp, J. Chem. Phys. 147, 152722 (2017).

[73] A. P. Lyubartsev, A. A. Martsinovski, S. V. Shevkunov, and P. N. Vorontsov-Velyaminov, J. Chem. Phys. 96, 1776 (1992).

[74] Z. Tan, J. Comput. Graph. Stat. 26, 54 (2017).

[75] A. C. Newton, J. Groenewold, W. K. Kegel, and P. G. Bolhuis, Proc. Nat. Acad. Sci. USA 112, 15308 (2015).

[76] P. Eastman, M. S. Friedrichs, J. D. Chodera, R. J. Radmer, C. M. Bruns, J. P. Ku, K. A. Beauchamp, T. J. Lane, L.-P. Wang, D. Shukla, T. Tye, M. Houston, T. Stich, C. Klein, M. R. Shirts, and V. S. Pande, J. Chem. Theory Comput. 9, 461 (2012).

[77] D. van der Spoel, E. Lindahl, B. Hess, G. Groenhof, A. E. Mark, and H. J. C. Berendsen, J. Comput. Chem. 26, 1701 (2005).

[78] B. Hess, C. Kutzner, D. van der Spoel, and E. Lindahl, J. Comput. Theory Chem. 4, 435 (2008).

[79] S. Plimpton, Journal of Computational Physics 117, 1 (1995).

[80] G. A. Tribello, M. Bonomi, D. Branduardi, C. Camilloni, and G. Bussi, Computer Physics Communications 185, 604 (2014).

[81] F. Noé and C. Clementi, Current Opinion in Structural Biology 43, 141 (2017).

[82] R. T. McGibbon, K. A. Beauchamp, M. P. Harrigan, C. Klein, J. M. Swails, C. X. Hernández, C. R. Schwantes, L.-P. Wang, T. J. Lane, and V. S. Pande, Biophysical Journal 109, 1528 (2015).

[83] K. A. Beauchamp, G. R. Bowman, T. J. Lane, L. Maibaum, I. S. Haque, and V. S. Pande, J. Chem. Theory Comput. 7, 3412 (2011).

[84] M. P. Harrigan, M. M. Sultan, C. X. Hernández, B. E. Husic, P. Eastman, C. R. Schwantes, K. A. Beauchamp, R. T. McGibbon, and V. S. Pande, Biophysical Journal 112, 10 (2017).

[85] M. K. Scherer, B. Trendelkamp-Schroer, F. Paul, G. Perez-Hernandez, M. Hoffmann, N. Plattner, C. Wehmeyer, J.-H. Prinz, and F. Noé, J. Chem. Theory Comput. 11, 5525 (2015).

[86] W. L. Jorgensen, J. Chandrasekhar, J. D. Madura, R. W. Impey, and M. L. Klein, The Journal of Chemical Physics 79, 926 (1983).

[87] P. A. Kollman, Accounts of Chemical Research 29, 461 (1996).

[88] J. D. Chodera, W.C. Swope, J.W. Pitera,and K.A. Dill, Multiscale Modeling& Simulation 5, 1214 (2006).

[89] J.-H. Prinz, J. D. Chodera, V. S. Pande, W. C. Swope, J. C. Smith, and F. Noé, J. Chem. Phys. 134, 244108 (2011).

[90] D. A. Sivak, J. D. Chodera, and G. E. Crooks, J. Phys. Chem. B 118, 6466 (2014).

[91] P. G. Bolhuis, C. Dellago, and D. Chandler, Proc. Natl. Acad. Sci. 97, 5877 (2000).

[92] P. G. Bolhuis and C. Dellago, The European Physical Journal Special Topics 224, 2409 (2015).

[93] H. Nguyen, D. A. Case, and A. S. Rose, Bioinformatics btx789 (2017).

[94] B. Leimkuhler and C. Matthews, J. Chem. Phys. 138, 174102 (2013).

[95] M. Shirts, Science 290, 1903 (2000).

[96] K. Lindorff-Larsen, S. Piana, R. O. Dror, and D. E. Shaw, Science 334, 517 (2011).

[97] B. Isralewitz, M. Gao, and K. Schulten, CurrentOpinion in Structural Biology 11, 224(2001).

[98] G. Henkelman and H. Jónsson, The Journal of Chemical Physics 113, 9978 (2000).

[99] W. e, W. Ren, and E. Vanden-Eijnden, The Journal of Physical Chemistry B 109, 6688 (2005).

